# Cd59 and inflammation orchestrate Schwann cell development

**DOI:** 10.1101/2022.01.11.475853

**Authors:** Ashtyn T. Wiltbank, Emma R. Steinson, Stacey J. Criswell, Melanie Piller, Sarah Kucenas

## Abstract

Efficient neurotransmission is essential for organism survival and is enhanced by myelination. However, the genes that regulate myelin and myelinating glial cell development have not been fully characterized. Data from our lab and others demonstrates that *cd59*, which encodes for a small GPI-anchored glycoprotein, is highly expressed in developing zebrafish, rodent, and human oligodendrocytes (OLs) and Schwann cells (SCs), and that patients with CD59 dysfunction develop neurological dysfunction during early childhood. Yet, the function of CD59 in the developing nervous system is currently undefined. In this study, we demonstrate that *cd59* is expressed in a subset of developing SCs. Using *cd59* mutant zebrafish, we show that developing SCs proliferate excessively, which leads to reduced myelin volume, altered myelin ultrastructure, and perturbed node of Ranvier assembly. Finally, we demonstrate that complement activity is elevated in *cd59* mutants and that inhibiting inflammation restores SC proliferation, myelin volume, and nodes of Ranvier to wildtype levels. Together, this work identifies Cd59 and developmental inflammation as key players in myelinating glial cell development, highlighting the collaboration between glia and the innate immune system to ensure normal neural development.

## Introduction

Myelin is a highly specialized, lipid-rich membrane that insulates axons in the nervous system to enhance neurotransmission (Ritchie, 1982) while preserving axon health (Stadelmann et al., 2019). Myelin is produced and maintained by myelinating glial cells such as motor exit point glia (Fontenas and Kucenas, 2021, 2018; Smith et al., 2014) and myelinating Schwann cells (SCs; Figure 1A) (Ackerman and Monk, 2016; Ben Geren, 1954; Jessen and Mirsky, 2005) in the peripheral nervous system (PNS) as well as oligodendrocytes (OLs) in the central nervous system (CNS) (Figure 1B) (Ackerman and Monk, 2016; Maturana, 1960; Nave and Werner, 2014; Peters, 1960a, 1960b). Normal myelin and myelinating glial cell development facilitate an efficient and effective nervous system, ensuring precise motor, sensory, and cognitive function (Almeida and Lyons, 2017; Berger et al., 2006; Stadelmann et al., 2019; Wei et al., 2019). When this process is impaired, as seen with inherited disorders that cause abnormal myelination, patients present with neurological dysfunction that often leads to severe physical and intellectual disabilities (Berger et al., 2006; Stadelmann et al., 2019; van der Knaap and Bugiani, 2017; Wei et al., 2019).

**Figure 1.**
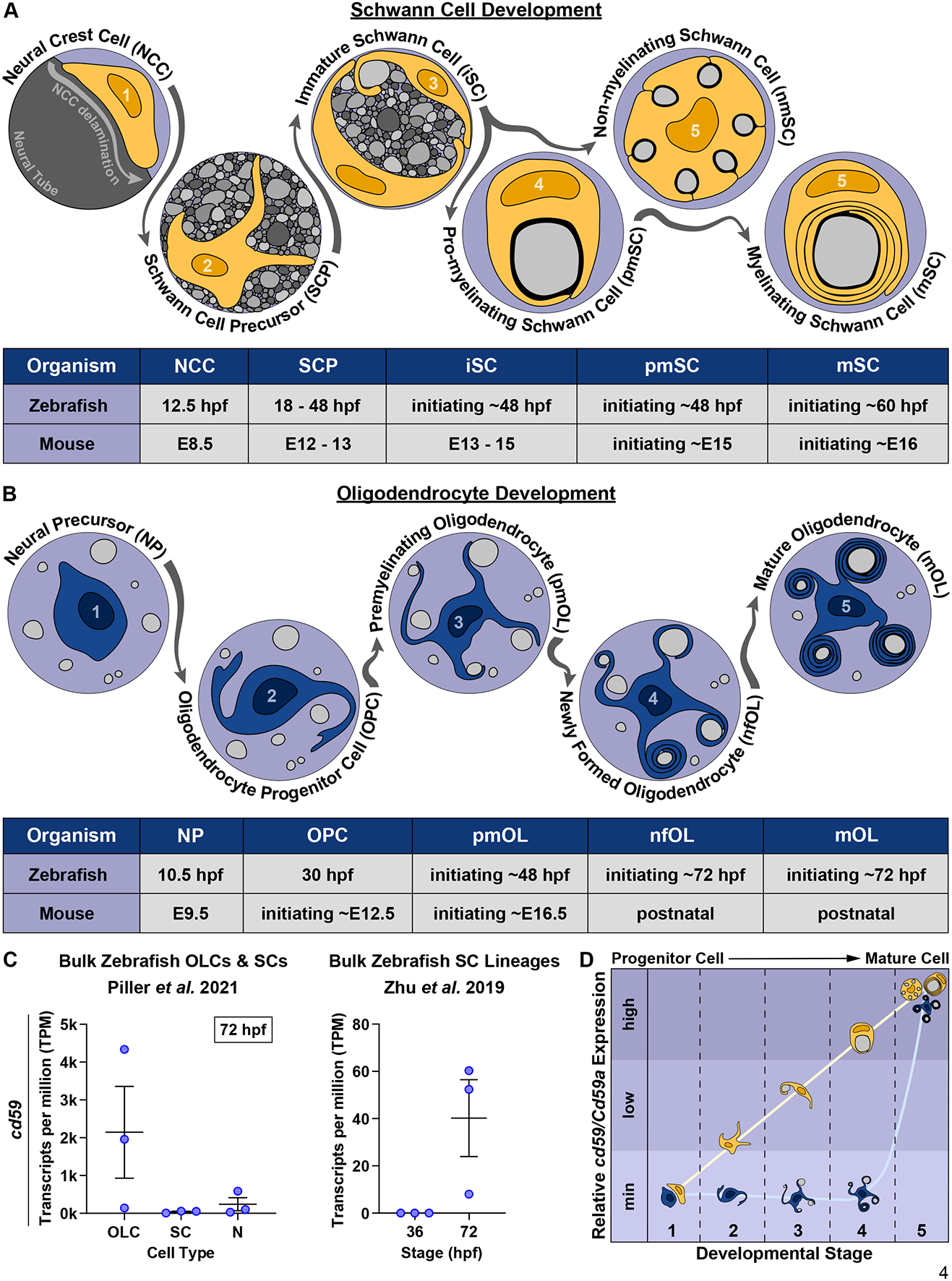
*cd59* is expressed in myelinating glial cells during nervous system development. (**A**) Timeline of SC (orange) development (top panel). SC developmental stages for zebrafish (hpf) and mice [embryonic day (E)] are indicated in the bottom panel. (**B**) Timeline of OL (blue) development. OL developmental stages for zebrafish (hpf) and mice (E and postnatal day) are indicated in the bottom panel. (**C**) Scatter plot of *cd59* expression (TPM) in OLCs, SCs, and neurons (N) at 72 hpf (left; mean ± SEM: OLC: 2145.1 ± 1215.1, SC: 40.1 ± 16.3; N: 240.5 ± 173.3; dot = replicate) as well as SCs at 36 and 72 hpf (right; mean ± SEM: 36 hpf: 0.0 ± 0.0, 72 hpf: 40.2 ± 16.3; dot = replicate). (**D**) Schematic of the relative *cd59/Cd59a* expression in developing SCs (orange) and OLs (blue) determined from RNAseq analysis in **Figure 1 - figure supplement 1**. Developmental stage numbers correspond with stages indicated in (**A**) and (**B**). Artwork created by Ashtyn T. Wiltbank with Illustrator (Adobe) based on previous schematics and electron micrographs published in (Ackerman and Monk, 2016; Cunningham and Monk, 2018; Jessen and Mirsky, 2005). **<u>Supplemental and Source Data for Figure 1</u>** **Figure 1 – figure supplement 1.** Myelinating glial cell *cd59/Cd59a* expression from bulk and scRNAseq. **Figure 1 – source data 1.** Source data for cd59 bulk, RNAseq expression.

Despite the importance of myelin, we still lack a complete understanding of myelin and myelinating glial cell development. With the recent boon of transcriptomic and proteomic analyses, many genes and proteins have been highlighted as differentially expressed during myelinating glial cell development, yet their precise functions remain unknown. For example, *cd59*, a gene that encodes for a small, glycosylphosphotidylinositol (GPI)- anchored glycoprotein, is a particularly interesting candidate for exploration. This gene features in several RNA sequencing (RNAseq) and proteomic analyses of zebrafish and rodent myelinating glial cells (Gerber et al., 2021; Howard et al., 2021; Marisca et al., 2020; Marques et al., 2018, 2016; Piller et al., 2021; Saunders et al., 2019; Siems et al., 2021; Wolbert et al., 2020; Zhu et al., 2019), all of which demonstrate high expression of *cd59* in developing oligodendrocyte lineage cells (OLCs) and SCs. *cd59* continues to be expressed in juvenile and adult zebrafish (Saunders et al., 2019; Siems et al., 2021; Sun et al., 2013) and becomes the 4^th^ most abundant CNS myelin protein in adult zebrafish (Siems et al., 2021). Phylogenetic analysis demonstrates that Cd59 is conserved within vertebrate evolution (Siems et al., 2021), and accordingly, CD59 protein is also found in developing and adult human myelin (Erne et al., 2002; Koski et al., 2002; Scolding et al., 1998; Zajicek, 1995). Collectively, this data indicates that *cd59* is a key component of the genetic program that orchestrates myelinating glial cell development. However, despite this knowledge of Cd59 expression, little is known about the function of Cd59 in the developing nervous system.

Outside of the nervous system, Cd59 is best known as a complement inhibitory protein. Complement is a molecular cascade within the innate immune system that aids in cell lysis of invading pathogens or aberrant cells in the body (Merle et al., 2015). This process lacks specificity and requires inhibitory proteins, such as Cd59, to protect healthy cells from lytic death (Davies et al., 1989). Cd59, specifically, acts at the end of the complement cascade where it binds to complement proteins 8 and 9 (C8 and C9, respectively) to prevent the polymerization of C9 and the subsequent formation of lytic pores (also known as membrane attack complexes (MACs)) in healthy cell membranes (Meri et al., 1990; Ninomiya and Sims, 1992; Rollins et al., 1991). This interaction is important throughout the adult body, including the nervous system, where CD59a is neuroprotective in models of multiple sclerosis (Mead et al., 2004) and neuromyelitis optica (Yao and Verkman, 2017). Beyond complement-inhibitory functions, Cd59 can also facilitate vesicle signaling in insulin-producing pancreatic cells (Golec et al., 2019; Krus et al., 2014), suppress cell proliferation in T cells responding to a viral infection (Longhi et al., 2005) or smooth muscle cells in models of atherosclerosis (Li et al., 2013), and orchestrate proximal-distal cell identity during limb regeneration (Echeverri and Tanaka, 2005). Like complement- inhibition, these processes are critical to normal nervous system function (Almeida et al., 2021; Baron and Hoekstra, 2010; Reiter and Bongarzone, 2020; White and Krämer-Albers, 2014). However, it is unclear exactly what role Cd59 is playing in myelinating glial cell development.

Though the precise function is unknown, it is clear that CD59 does impact human nervous system development. Patients with CD59 dysfunction, such as those with congenital CD59 deficiency (Haliloglu et al., 2015; Höchsmann and Schrezenmeier, 2015; Karbian et al., 2018; Solmaz et al., 2020) or germline paroxysmal nocturnal hemoglobinuria (Johnston et al., 2012), present with polyneuropathies during infancy and continue to have nervous system dysfunction throughout their lives. Intriguingly, these neurological symptoms persist even with complement inhibition, the most common treatment for congenital CD59 deficiency (Höchsmann and Schrezenmeier, 2015). Together, these observations further indicate that CD59 has an additional role in the developing nervous system and requires further investigation.

Here, utilizing the zebrafish model, we examined the role of Cd59 in the developing nervous system. In this study, we found that *cd59* is highly expressed in a subset of developing SCs in addition to mature OLs and myelinating and non-myelinating SCs. We chose to focus on SCs and found that global mutation of *cd59* resulted in excessive proliferation of developing SCs as well as impaired myelin and node of Ranvier development. Finally, we demonstrate that complement activity increases in *cd59* mutants and that unregulated inflammation contributes to SC overproliferation and aberrant myelin and node of Ranvier formation. Overall, this data demonstrates that developmental inflammation stimulates SC proliferation and that this process is balanced by Cd59 to ensure normal SC and myelinated nerve development.

## Results

### *cd59* is expressed in myelinating glial cells during nervous system development

Myelinating glial cell development is a complex process (Figure 1A, B) orchestrated by a genetic program that is not yet fully elucidated. To identify unexplored genes that may impact myelinating glial cell or myelin development, we evaluated bulk and single-cell (sc) RNAseq datasets that assessed myelinating glial cells during nervous system development (Gerber et al., 2021; Howard et al., 2021; Marisca et al., 2020; Marques et al., 2018, 2016; Piller et al., 2021; Saunders et al., 2019; Wolbert et al., 2020; Zhu et al., 2019). Across multiple datasets, *cd59* (zebrafish) or *Cd59a* (mouse), which encodes for a small, GPI-anchored glycoprotein that has no known function in the developing nervous system, is expressed in SCs and OLCs (Figure 1C and Figure 1 – figure supplement 1).

When looking at SCs, data from developing zebrafish and mice indicated that *cd59/Cd59a*, respectively, is minimally expressed in neural crest cells (NCCs; the multipotent progenitors that give rise to SCs; Figure 1 – figure supplement 1A) (Howard et al., 2021; Zhu et al., 2019) but increases in expression in SC precursors (SCPs; Figure 1 – figure supplement 1B.1, B.2) (Gerber et al., 2021), immature SCs (iSCs; Figure 1 – figure supplement 1B.1, B.2) (Gerber et al., 2021), pro-myelinating SCs (pmSCs; Figure 1 – figure supplement 1B.1, B.2) (Gerber et al., 2021), and mature myelinating and non-myelinating SCs (MSCs and NMSCs; Figure 1C and Figure 1 – figure supplement 1B.3, C.1, C.2) (Gerber et al., 2021; Piller et al., 2021; Saunders et al., 2019; Wolbert et al., 2020). This data indicates that Cd59 may be important during early stages of SC development as well as in mature MSCs and NMSCs (Figure 1D).

In contrast, bulk RNAseq of zebrafish oligodendrocyte progenitor cells (OPCs) and OLs indicated that *cd59* is expressed near the onset of myelination (72 hpf; Figure 1C) (Piller et al., 2021). scRNAseq (Marisca et al., 2020; Marques et al., 2018, 2016) and *in situ* hybridization (ISH) (Siems et al., 2021) of zebrafish and mouse OLCs showed that *cd59/CD59a* is mostly expressed in mature OLs and not OPCs (Figure 1 – figure supplement 1D). CD59 protein is also present within newly-formed human myelin sheaths as is evident in immunostaining of third trimester fetal brains (Zajicek, 1995). Based on these findings, it is likely that Cd59 does not play a role in early stages of OLC development but could influence mature, myelinating OL function (Figure 1D).

Intrigued by this elevated expression of *cd59* in developing myelinating glial cells, we verified that *cd59* RNA was expressed in OLCs and SCs with chromogenic and fluorescent ISH (CISH and FISH, respectively) at 3 dpf (Figure 2A, B). Morphology, location, and co-expression with *sox10:megfp*, which is a marker for both OLCs and SCs, showed that *cd59* RNA is expressed in both cell types at 3 dpf (Figure 2A, B). Beyond expression in the heart and pancreas, *cd59* expression is largely confined to developing myelinating glial cells at this stage.

**Figure 2.**
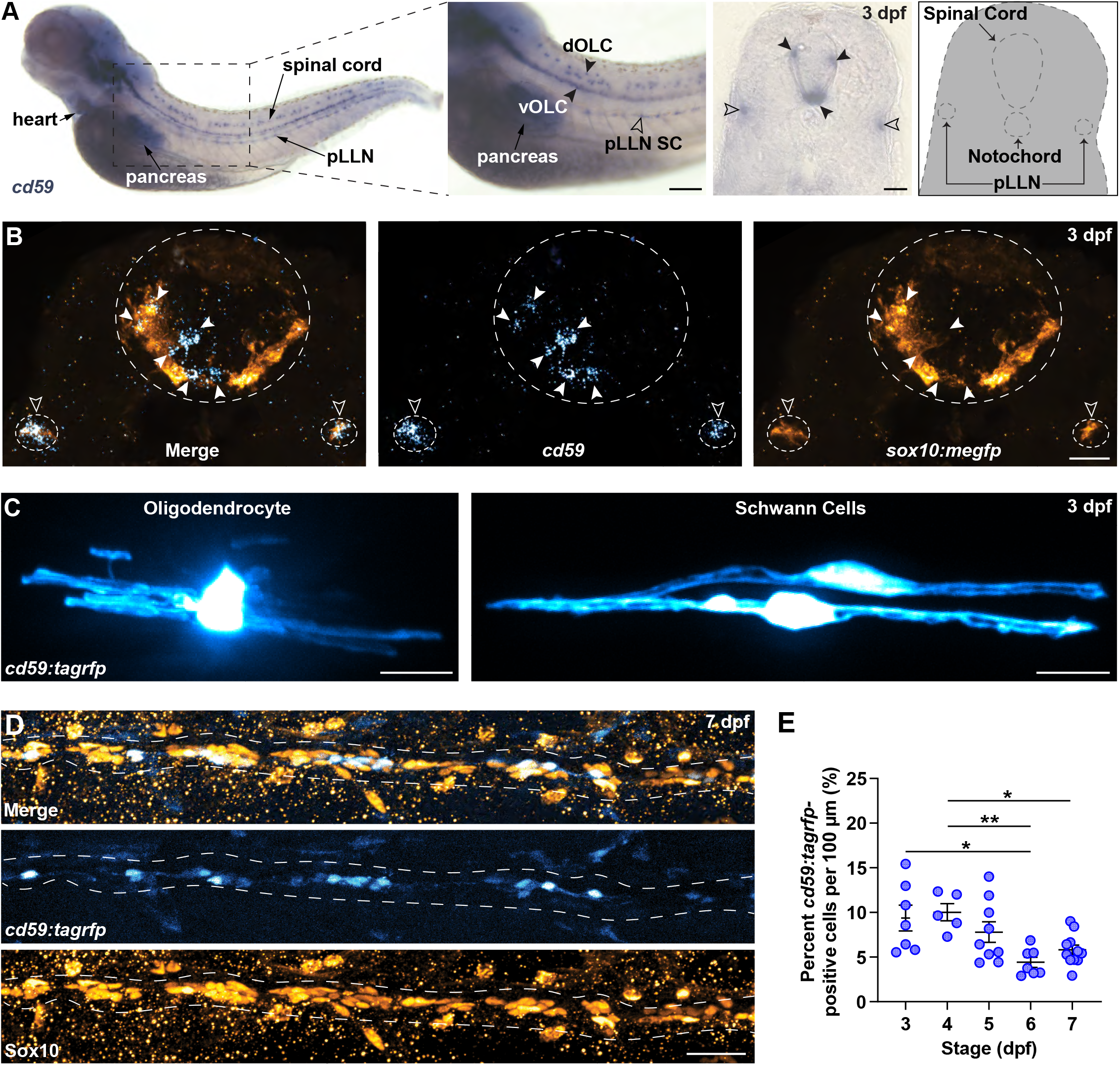
*cd59* is expressed in SCs and OLs. (**A**) Whole mount CISH showing *cd59* expression (purple) in the heart, pancreas, dorsal and ventral OLCs (dOLCs and vOLCs, respectively; filled arrows), and pLLN SCs (empty arrows) at 3 dpf. Schematic (right panel) indicates location of spinal cord, notochord, and pLLN in transverse section. (**B**) FISH showing *cd59* expression (cyan) in *sox10:megfp*-positive, pLLN SCs (orange cells indicated by empty arrows) and spinal cord OLs (orange cells indicated by filled arrows) at 3 dpf in transverse sections. (**C**) Mosaic labeling showing a *cd59:tagrfp*-positive OL in the spinal cord (left) and two SCs on the pLLN (right) at 3 dpf. (**D**) Immunofluorescence (IF) showing *cd59:tagrfp* expression (cyan) in Sox10-positive SCs (orange) along the pLLN at 7 dpf. White dashed lines outline the pLLN. Sox10-positive pigment cells outside of the dashed lines were not included in the analysis. (**E**) Scatter plot of percent *cd59:tagrfp*-positive SCs on the pLLN from 3 to 7 dpf (mean ± SEM: 3 dpf: 9.4 ± 1.5, 4 dpf: 10.0 ± 1.0, 5 dpf: 7.8 ± 1.2, 6 dpf: 4.4 ± 0.6, 7 dpf: 5.8 ± 0.6; p values: 3 dpf vs 6 dpf: p = 0.0126, 4 dpf vs 6 dpf: p = 0.0095, 4 dpf vs 7 dpf: p = 0.0477; dot = 1 fish). Data collected from somites 11 to 13 (∼320 µm) and normalized to units per 100 µm. Scale bars: (**A**) lateral view, 100 µm; transverse section, 25 µm; (**B, C**) 10 µm; (**D**) 25 µm. Artwork created by Ashtyn T. Wiltbank with Illustrator (Adobe). **<u>Supplemental and Source Data for Figure 2</u>** **Figure 2—Figure Supplement 1.** *cd59* is expressed in the floorplate, hypochord, and developing SCs. **Figure 2—Figure Supplement 2.** *cd59:tagrfp*-positive SCs express canonical SC markers. **Figure 2 – source data 1.** Source data for quantification of *cd59:tagrfp*-positive cells.

To study these cells *in vivo*, we created a transcriptional reporter zebrafish line for *cd59* (herein referred to as *cd59:tagrfp)*. This line utilizes 5 kilobases (kb) of the *cd59* promoter as well as the first exon and intron, the latter containing an enhancer that is important for *cd59* expression (Tone et al., 1999), to drive expression of TagRFP in *cd59*-expressing cells. With *in vivo,* confocal imaging, mosaic-labeling with our *cd59:tagrfp* construct revealed labeling of OLs and SCs with myelin-like processes (Figure 2C), indicating that *cd59* is expressed in mature myelinating glial cells.

Bulk and scRNAseq data indicated that *cd59* is expressed during early stages of SC development whereas *cd59* expression was restricted to mature OLs. To verify this expression pattern, we looked at *cd59* RNA expression at earlier stages in the SC lineage. At 24 hpf, using CISH and FISH, we did not observe *cd59* expression in *sox10:megfp*-positive NCCs, though we did see it in the hypochord and floor plate of the spinal cord at this stage (Figure 2 – figure supplement 1A, B). At 36 and 48 hpf, when NCCs begin to develop into SCPs and iSCs, we detected sporadic expression of *cd59* with FISH in *sox10:megfp*-positive cells along the posterior lateral line nerve (pLLN) in the PNS (Figure 2 – figure supplement 1C). By 72 hpf, we observed robust *cd59* expression within *sox10:megfp-*positive SCs along the pLLN (Figure 2 – figure supplement 1C). These data demonstrate that *cd59* is expressed in developing SCs, confirming the expression patterns seen in the RNAseq datasets we analyzed (Figure 1 – figure supplement 1).

From our ISH data, we noticed that *cd59* was not expressed in every SC. Notably, SCs associated with spinal motor nerves did not express *cd59* (Figure 2A, B). Rather, *cd59* expression was largely confined to SCs along the pLLN (Figure 2A, B). To investigate this expression further, we utilized our stable *cd59:tagrfp* line. This transgenic line provides clear, *in vivo* labeling of c*d59*-expressing pLLN SCs, which we confirmed using FISH with a *cd59* probe in *cd59:tagrfp-*positive SCs (Figure 2 – figure supplement 2A). To verify that *cd59:tagrfp*-positive SCs expressed canonical SC markers, including *SRY-box transcription factor 10* (*sox10)*, *erb-b2 receptor tyrosine kinase 3b* (*erbb3b)*, and *myelin basic protein a* (*mbpa)*, we examined pLLN SCs in transgenic and gene trap larvae co-expressing these markers with *in vivo* confocal imaging. As expected, *cd59:tagrfp* expression was observed in *sox10, erbb3b,* and *mbp-*positive SCs at 72 hpf and 7 dpf (Figure 2 – figure supplement 2B – D), confirming that *cd59:tagrfp-*positive SCs express other known SC genes.

To determine whether *cd59* was expressed in all pLLN SCs, we used immunofluorescence (IF) to co-label *cd59:tagrfp*-positive SCs with an antibody against Sox10, which labels all SCs along the pLLN. Strikingly, when looking at SCs from 3 to 7 dpf, *cd59:tagrfp* was only expressed in a subset of pLLN SCs (average of 4.4 ± 1.5% to 9.4 ± 3.8% of pLLN SCs; Figure 2D, E). We also noted that *cd59* expression was only expressed in a subset of OLs (Figure 2B) and was completely lacking in MEP glia. Taken together, these data demonstrate that *cd59* expression is heterogeneous in developing myelinating glial cells.

### Generation of *cd59* mutant zebrafish with CRISPR/Cas9 genome editing

Our findings demonstrate that *cd59* is expressed in a subset of developing SCs. However, it is unclear how Cd59 influences SC development. In other cells, Cd59 has many different functions, including preventing complement-dependent cell lysis (Davies et al., 1989), facilitating vesicle-dependent signaling (Golec et al., 2019; Krus et al., 2014), suppressing cell proliferation (Dashiell et al., 2000; Hila et al., 2001; Li et al., 2013; Longhi et al., 2005), and orchestrating proximal-distal cell identity (Echeverri and Tanaka, 2005). There is little known about Cd59 in the developing nervous system, however.

To evaluate the function of Cd59 in SC development, we generated *cd59* mutant zebrafish with CRISPR/Cas9 genome editing. Targeting the end of the first coding exon (exon 2; Figure 3A), we recovered an allele with a 15 bp deletion that occurred across the exon 2 – intron 2 splice site, generating a splice mutation (herein referred to as mutant *cd59^uva48^*; Figure 3B). RT-PCR and Sanger sequencing analysis of mutant *cd59^uva48^* transcripts revealed that multiple splice variants were produced compared to wildtype *cd59*, which produced a single transcript (Figure 3 – figure supplement 1A, E). Analysis of the mutant transcripts indicated that an early stop codon was generated in the N-terminal signal sequence, producing a severely truncated protein (10 – 48 AA compared to the 119 AA wildtype protein; Figure 3 – figure supplement 1E – G). Furthermore, the predicted proteins lacked the Ly6/uPAR domain and GPI-anchor signal sequence (Figure 3 – figure supplement 1E – G). Therefore, the mutant Cd59 protein was expected to be non-functional. We also recovered an additional *cd59* mutant allele, *cd59^uva47^*, which was produced with the same single guide RNA (sgRNA). This mutant had a 6 bp deletion near the end of exon 2 that did not affect the splice site (Figure 3 – figure supplement 1B, D) and therefore produced a single mutant transcript (Figure 3 – figure supplement 1B). Sequencing of the mutant *cd59^uva47^* transcript predicted the absence of two amino acids (AA) in the N-terminal signal sequence, which would yield a slightly smaller protein (177 AA) compared to wildtype Cd59 (119 AA; Figure 3 – figure supplement 1E – G). Though the protein was not severely truncated, we predicted that interfering with the N-terminal signal sequence would prevent protein trafficking to the endoplasmic reticulum and interrupt proper protein function. Collectively, mutant *cd59^uva48^* larvae have perturbed RNA splicing that results in premature termination codon (PTC) formation and probable protein truncation, and mutant *cd59^uva47^* larvae have a disrupted N-terminal sequence. Considering these results, both mutants were suspected to have impaired Cd59 function.

**Figure 3.**
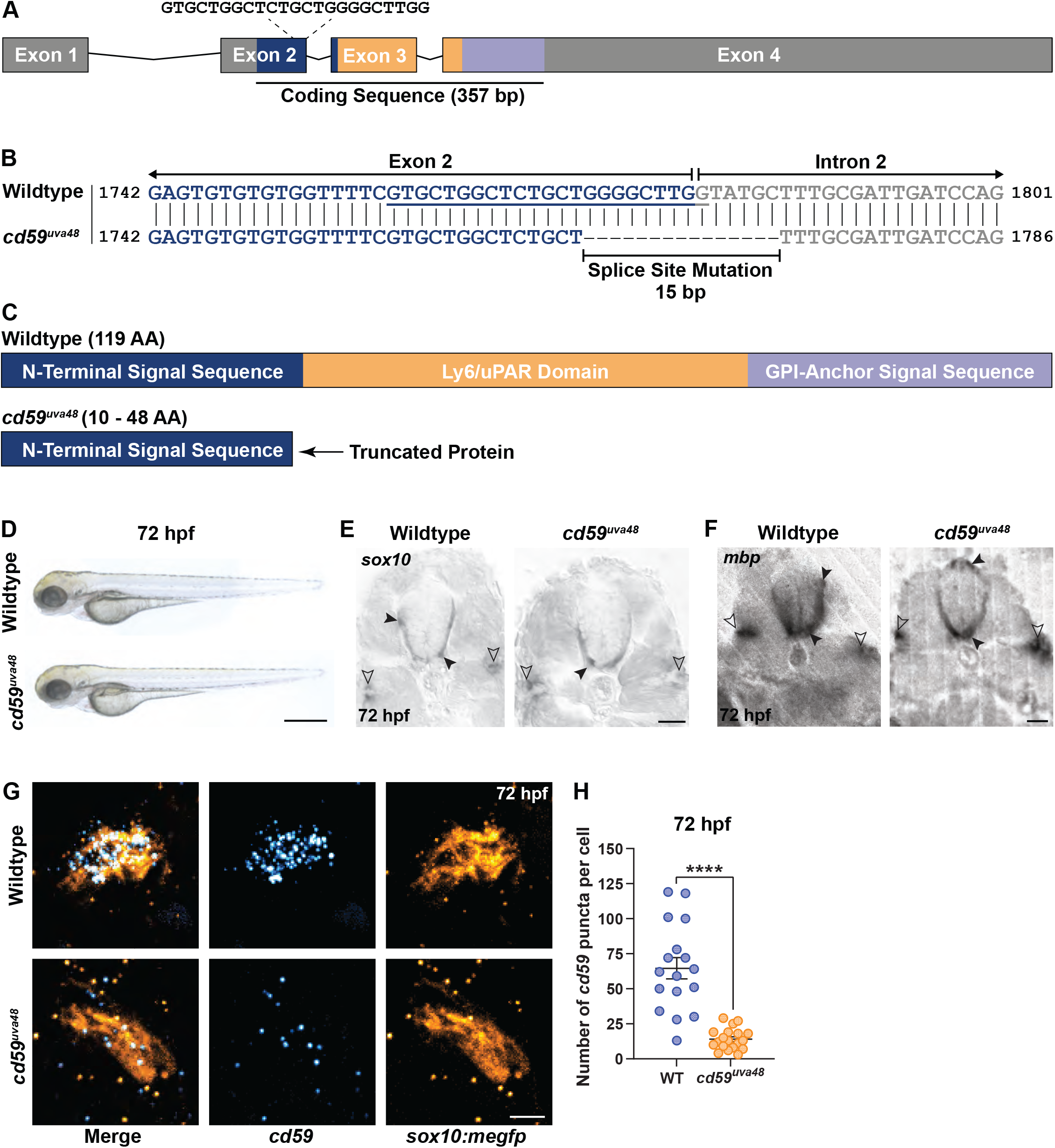
Generation of *cd59* mutant zebrafish with CRISPR/Cas9 genome editing. **(A)** Schematic of zebrafish *cd59* gene. The *cd59* coding sequence (357 bp; CDS) encodes the protein domains with the corresponding colors indicated in **(C)**. The non-CDS is indicated in grey. The dashed lines indicate the sgRNA and the target sequence. **(B)** Genomic sequences for wildtype *cd59* and mutant *cd59^uva48^* showing a 15 bp deletion at the splice site between exon 2 (blue) and intron 2 (grey) of the *cd59* gene. (**C**) Schematic of Cd59 protein made in wildtype (119 AA; top panel) and *cd59^uva48^* mutant fish (10 to 48 AA; bottom panel). (**D**) Bright-field images of wildtype and *cd59^uva48^* mutant larvae at 72 hpf showing no anatomical defects as a result of *cd59* mutation. (**E**) CISH showing *sox10*-positive (grey) OLs in the spinal cord (filled arrows) and SCs on the pLLN (empty arrows) at 72 hpf in transverse sections. (**F**) CISH showing *mbp*-positive (grey) OLs in the spinal cord (filled arrows) and SCs on the pLLN (empty arrows) at 72 hpf in transverse sections. (**G**) FISH showing *cd59* expression (cyan) in *sox10:megfp*-positive SCs (orange) along the pLLN at 72 hpf. (**H**) Scatter plot of the number of *cd59* RNA puncta in pLLN SCs (mean ± SEM: WT: 64.7 ± 7.6, *cd59^uva48^*: 14 ± 1.7; p < 0.0001; dot = 1 cell; n = 7 fish per group). Scale bars: (**A**) 0.25 mm; (**E, F**) 25 µm; (**G**) 5 µm. Artwork created by Ashtyn T. Wiltbank with Illustrator (Adobe). **<u>Supplemental and Source Data for Figure 3</u>** **Figure 3 - figure supplement 1.** Characterization of *cd59* mutant zebrafish. **Figure 3 – source data 1.** Source data from quantification of *cd59* puncta. **Figure 3 – figure supplement 1 – source data 1.** Source data from gel electrophoresis of RT-PCR of *cd59^uva48^* mutant embryos. **Figure 3 – figure supplement 1 – source data 2.** Source data from gel electrophoresis of RT-PCR of *cd59^uva47^* mutant embryos.

To begin our initial evaluation of these mutants, we sought to compare general developmental characteristics between *cd59* mutant larvae and their wildtype siblings. Both *cd59^uva48^* and *cd59^uva47^* homozygous adults were viable and produced embryos with no anatomical, behavioral, or reproductive defects (Figure 3D and Figure 3 – figure supplement 1C). Next, we wanted to know if SCs were present in mutant *cd59^uva48^* larvae, which were utilized in the majority of our studies. Using CISH, we labeled wildtype and mutant *cd59^uva48^* larvae with RNA probes against *sox10* and *mbp* at 72 hpf. Both wildtype and mutant larvae had *sox10* and *mbp-*positive SCs along the pLLN (Figure 3E, F) indicating that Cd59 was not necessary for SC genesis.

Knowing that SCs were present, we were then curious how *cd59* expression was affected in *cd59^uva48^* mutant SCs. According to our RT-PCR analysis, mutant *cd59^uva48^* embryos produced RNA transcripts with PTCs (Figure 3 – figure supplement 1E). RNA transcripts with PTCs can sometimes undergo nonsense-mediated decay (NMD), a process by which normal and mutant gene expression is controlled through RNA degradation (Bruno et al., 2011; Chang et al., 2007; Karam and Wilkinson, 2012; Lou et al., 2014). To determine if mutant *cd59^uva48^* RNA transcripts underwent NMD, we used FISH to label wildtype and mutant *cd59^uva48^;sox10:megfp* larvae with a probe against *cd59* (Figure 3G). Quantification of *cd59* RNA puncta showed that *cd59* expression was significantly reduced in mutant SCs compared to wildtype SCs (Figure 3H), indicating that *cd59^uva48^* mutant RNA was being degraded through NMD and therefore would result in a loss of protein. Overall, these findings demonstrate that *cd59^uva48^* mutants can be used as a loss-of-function model in which to study the role of Cd59 in developing SCs.

### Cd59 regulates developing Schwann cell proliferation

With our new *cd59* mutants, we began to explore the possible roles Cd59 could play in SC development. Previous studies demonstrate that CD59a can prevent overproliferation of T cells during a viral infection (Longhi et al., 2005). Similarly, CD59a is important for limiting deleterious smooth muscle cell proliferation during atherosclerosis (Li et al., 2013). Because we observed *cd59* expression during the proliferative phases of SC development, we hypothesized that Cd59 could be involved in developmental SC proliferation. To test our hypothesis, we labeled SCs with a Sox10 antibody and quantified the number of Sox10-positive SCs along the pLLN in wildtype and *cd59^uva48^* mutant larvae at various stages during SC development (36 hpf to 7 dpf) (Figure 4A). At all stages we investigated, there were significantly more SCs in mutant embryos and larvae compared to wildtype siblings (Figure 4A). We observed this excess early in SC development during the SCP stage (36 hpf; Figure 4B) through the iSC and pmSC stages (∼48 hpf, Figure 4C) and into mature MSC stages (72 hpf, Figure 4D; 7 dpf, Figure 4E) (Ackerman and Monk, 2016). To rule out the possibility of off-target effects from CRISPR/Cas9 genome editing, we evaluated *cd59^uva47^* mutants as well. Similarly, we observed the same phenotype in mutant *cd59^uva47^* embryos at 48 hpf (Figure 4 – figure supplement 1A, B) demonstrating that mutation of *cd59* was responsible for this phenotype. Together, these data demonstrate that Cd59 functions to limit overproliferation of SCs during development.

**Figure 4.**
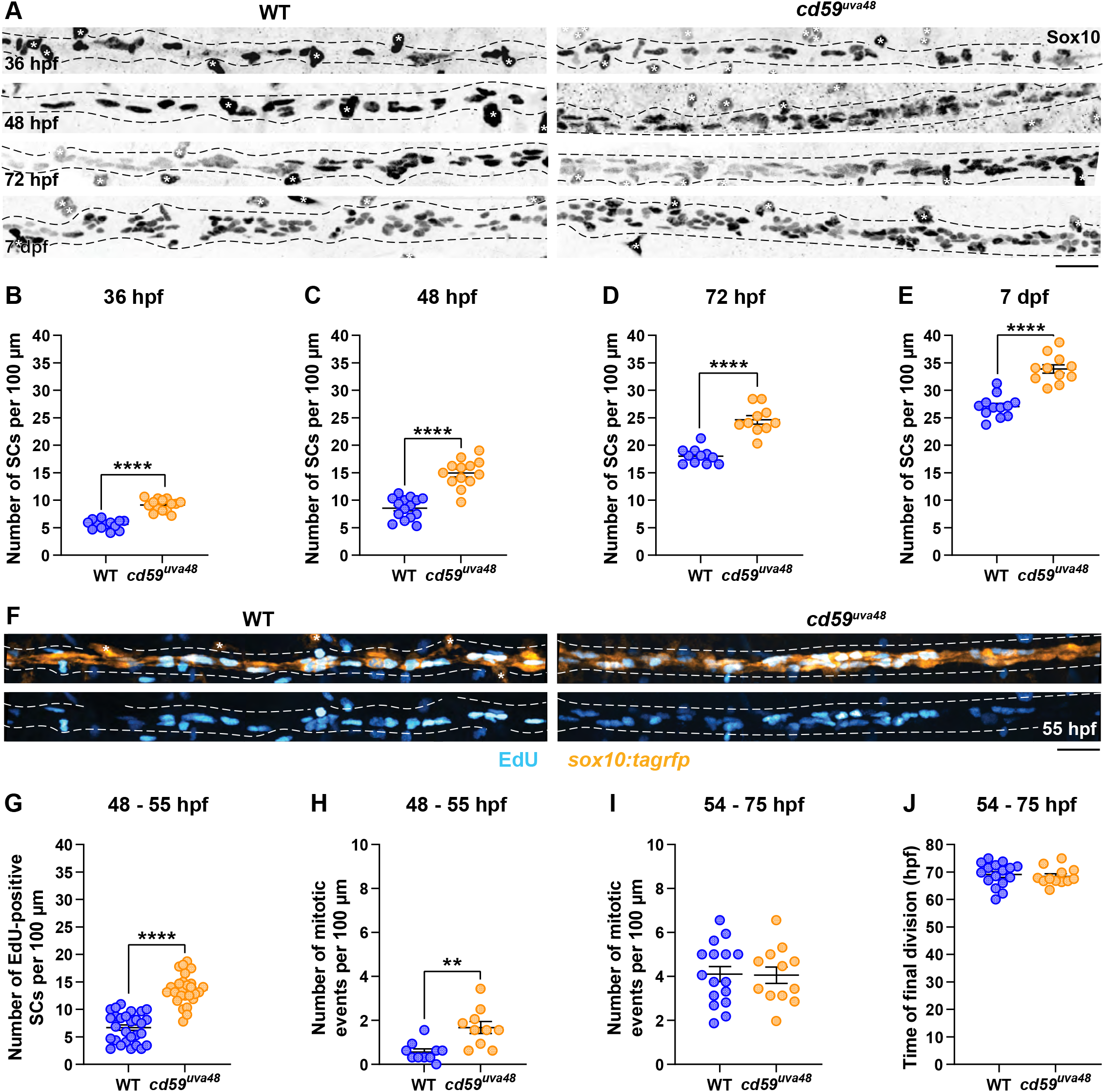
*cd59* regulates SC proliferation. (**A**) IF showing Sox10-positive SCs (black/grey) along the pLLN from 36 hpf to 7 dpf. Black dashed lines outline the pLLN. Sox10-positive pigment cells outside of the dashed lines (white asterisks) were not included in the analysis. (**B - E**) Scatter plots of the number of Sox10-positive SCs along the pLLN from 36 hpf to 7 dpf (mean ± SEM: 36 hpf: WT: 5.6 ± 0.2, *cd59^uva48^*: 9.1 ± 0.3; 48 hpf: WT: 8.6 ± 0.5, *cd59^uva48^*: 15.0 ± 0.7; 72 hpf: WT: 18.0 ± 0.4, *cd59^uva48^*: 24.3 ± 0.8; 7 dpf: WT: 27.0 ± 0.6, *cd59^uva48^*: 33.9 ± 0.8; p values: p < 0.0001; dot = 1 fish). (**F**) EdU incorporation assay showing *sox10:tagrfp*-positive, pLLN SCs (orange) pulsed with EdU (cyan) from 48 to 55 hpf. (**G**) Scatter plot of the number of EdU-positive SCs along the pLLN at 55 hpf (WT: 6.7 ± 0.5, *cd59^uva48^*: 13.6 ± 0.5; p < 0.0001; dot = 1 fish). (**H**) Scatter plot of the number of mitotic events observed in SCs from 48 to 55 hpf (mean ± SEM: WT: 0.6 ± 0.1, *cd59^uva48^*: 1.7 ± 0.3; p = 0.0019; dot = 1 fish). (**I**) Scatter plot of the number of mitotic events observed in SCs from 54 to 75 hpf (mean ± SEM: WT: 4.1 ± 0.3, *cd59^uva48^*: 4.1 ± 0.4; dot = 1 fish). (**H**) Scatter plot of the time of final cell division (hpf) observed in SCs from 54 to 75 hpf (mean ± SEM: WT: 69.1 ± 1.1, *cd59^uva48^*: 68.4 ± 0.9; dot = 1 fish). All data were collected from somites 11 to 13 (∼320 µm) and normalized to units per 100 µm. Scale bars: (**A, F**) 25 µm. **<u>Supplemental and Source Data for Figure 4</u>** **Figure 4—Figure Supplement 1.** *cd59* does not impact proliferation of neurons or NCCs. **Figure 4—Figure Supplement 2.** Loss of *cd59* does not impact cell death within the PNS and CNS. **Figure 4 – source data 1.** Source data from quantification of pLLN SCs at 36 hpf. **Figure 4 – source data 2.** Source data from quantification of pLLN SCs at 48 hpf. **Figure 4 – source data 3.** Source data from quantification of pLLN SCs at 72 hpf. **Figure 4 – source data 4.** Source data from quantification of pLLN SCs at 7 dpf. **Figure 4 – source data 5.** Source data from quantification of EdU-positive SCs from 48 to 55 hpf. **Figure 4 – source data 6.** Source data for quantification of mitotic events from 48 to 55 hpf. **Figure 4 – source data 7.** Source data for quantification of mitotic events from 54 to 75 hpf. **Figure 4 – source data 8.** Source data for time of final division. **Figure 4 – figure supplement 1 – source data 1.** Source data for quantification of SCs in *cd59^uva47^* mutant embryos. **Figure 4 – figure supplement 1 – source data 2.** Source data for quantification of pLLG neurons. **Figure 4 – figure supplement 1 – source data 3.** Source data for quantification of pLLG- associated NCCs. **Figure 4 – figure supplement 1 – source data 4.** Source data for quantification of migrating NCCs. **Figure 4 – figure supplement 1 – source data 5.** Source data for quantification of trunk neuromasts. **Figure 4 – figure supplement 2 – source data 1.** Source data for the quantification of AO-positive SCs. **Figure 4 – figure supplement 2 – source data 2.** Source data for the quantification of AO-positive cells ventral to the pLLN. **Figure 4 – figure supplement 2 – source data 3.** Source data for the quantification of TUNEL-positive SCs. **Figure 4 – figure supplement 2 – source data 4.** Source data for the quantification of TUNEL-positive CNS cells.

To independently confirm this excessive proliferation phenotype seen in SCs, we used EdU to label mitotically-active SCs during SC development. Specifically, we incubated *sox10:tagrfp* embryos in EdU from 48 to 55 hpf. The embryos were then fixed and the number of EdU-positive SCs on the pLLN were quantified with confocal imaging. As we hypothesized, the total number of EdU-positive SCs was increased in *cd59^uva48^* mutant embryos compared to wildtypes (Figure 4F & 4G). From these data, we confirmed that there was more SC proliferation occurring in *cd59^uva48^* mutant larvae when compared to wildtype larvae. Consistent with this finding, *in vivo,* time-lapse imaging of *sox10:tagrfp-*positive SCs along the pLLN showed *cd59^uva48^* mutant SCs undergoing significantly more mitotic events than their wildtype siblings, with most of the increased cell divisions occurring between 48 and 55 hpf (Figure 4H, Video 1, and Video 2). Between 54 and 75 hpf, however, the number of divisions was similar between wildtype and *cd59^uva48^* mutant larvae with no difference in the timing of terminal division (Figure 4I, J, Video 3, and Video 4), demonstrating that, although Schwann proliferation is initially amplified in *cd59^uva48^* mutant larvae, they are capable of terminating proliferation at the same developmental stage as wildtype larvae.

Taken together, these data demonstrate that Cd59 restricts developmental SC proliferation and that excess SCs generated in *cd59* mutant larvae persist past developmental stages. To rule out changes in cell death as a contributor to SC number, we used acridine orange (AO) incorporation and TUNEL staining to quantify the number of dying SCs at 48 hpf. Co-labeling of *sox10:tagrfp*-positive SCs with AO revealed no changes in SC death between wildtype and mutant *cd59^uva48^* embryos (Figure 4 – figure supplement 2A, B). When we assayed death more broadly in the trunk ventral to the pLLN, we also did not observe any increase in AO labeling, indicating that cell death in the embryo did not increase with *cd59* mutation (Figure 4 – figure supplement 2A, C). Similarly, TUNEL staining combined with Sox10 labeling at 48 hpf on sectioned tissue showed that apoptotic SC death on the pLLN was unaltered in mutant *cd59^uva48^* embryos (Figure 4 – figure supplement 2D, E). There was also no difference in the number of TUNEL-positive cells in the spinal cord (Figure 4 – figure supplement 2D, F). Finally, in agreement with this data, we observed no SC death during *in vivo* imaging of *sox10:tagrfp*-positive SCs from 48 to 75 hpf in both wildtype and *cd59^uva48^* larvae (Videos 1 - 4). Overall, these findings show that *cd59* mutation does not perturb SC death.

Finally, we wanted to determine if *cd59* mutation affected the proliferation of other neural cells in addition to SCs. Looking at CNS neurons labeled with a HuC/HuD antibody, we observed that spinal cord neuron density was indistinguishable between wildtype and mutant *cd59^uva48^* embryos at 48 hpf (Figure 4 – figure supplement 1C), supporting our observations that cell death does not change in the spinal cord (Figure 4 – figure supplement 2D, F). To determine if this was also the case in the PNS, we used the same HuC/HuD antibody to label posterior lateral line ganglia (pLLG) neurons and found that the number of HuC/HuD-positive pLLG neurons was the same between wildtype and *cd58^uva48^* mutant embryos at 24 hpf (Figure 4 – figure supplement 1D, E). These data indicate the neuronal proliferation is unaffected by *cd59* mutation. We also investigated whether NCC development was affected in our *cd59* mutant embryos. In accordance with the lack of *cd59* expression seen by RNAseq and ISH (Figure 1 – figure supplement 1A and Figure 2 – figure supplement 2A, B), we saw no difference in the number of cranial NCCs associated with the pLLG (Figure 4 – figure supplement 1D, F) nor in the number of migrating spinal motor nerve NCCs (Figure 4 – figure supplement 1G, H). Overall, these data show that Cd59 does not influence NCC proliferation. Furthermore, the increase in SC number during development was not due to an increase in the NCCs that give rise to SCs.

Another important aspect of the pLLN is that it innervates neuromasts, which are sensory organs that detect water movement (Ghysen and Dambly-Chaudière, 2007). Trunk neuromast number and positioning changes as the zebrafish grows in length, initially starting with 5 to 6 primary neuromasts (deposited from 20 to 40 hpf) that eventually expand into nearly 600 neuromasts in adults (Ghysen and Dambly-Chaudière, 2007). Developing SCs can impact the number of neuromasts, as is evident by inappropriate neuromast formation in *erbb* pathway and *sox10* mutants which lack SCs (Grant et al., 2005; Lush and Piotrowski, 2014; Perlin et al., 2011; Rojas-Muñoz et al., 2009). To determine if excess SCs also affect neuromast formation, we labeled neuromasts with an acetylated α-Tubulin antibody and quantified the number of truncal neuromasts. Interestingly, there was no change in truncal neuromast number when comparing wildtype and *cd59^uva48^* mutant larvae at 7 dpf (Figure 4 – figure supplement 1I), indicating that an increase in SC number does not alter neuromast formation. Combined, these data demonstrate that Cd59 is playing an early role in SC development by limiting proliferation and that this proliferative effect does not extend to other neural cells in the CNS or PNS.

### Cd59 is necessary for myelin and node of Ranvier formation in the PNS

From our observations, we saw that Cd59 functions to prevent overproliferation of developing SCs. Therefore, we next wanted to understand how an increase in SC number would impact myelin and pLLN development. Proliferation is a critical aspect of radial sorting, the process by which iSCs segregate large caliber axons that require myelination, and is necessary for nerve development (Feltri et al., 2016). When proliferation is dysregulated, either through insufficient or excessive Schwann cell division, radial sorting can be delayed, arrested, or improperly executed, resulting in myelination that is incomplete or inappropriate, such as instances of polyaxonal myelination of small caliber axons (Feltri et al., 2016; Gomez-Sanchez et al., 2009). Therefore, we were curious how limiting SC proliferation through Cd59 influences myelinated nerve development. ISH for *mbp* (Figure 3F), confocal imaging of *mbpa:tagrfp-caax*-positive pLLN SCs (Figure 5 – figure supplement 1A – C), and transmission electron microscopy (TEM) (Figure 5A) showed that *cd59^uva48^* mutant SCs were capable of producing myelin. Additionally, mosaic myelin labeling with *mbpa:tagrfp-caax* or *mbpa:tagrfp-caax* and *mbp:egfp-caax* constructs, injected into one-cell embryos, indicated that the myelin sheath length at 7 dpf did not vary between wildtype and *cd59^uva48^* mutant larvae (Figure 5 – figure supplement 1A, D) nor was there any evidence of overlapping sheaths at the same age [data not shown]. However, when looking in *mbpa:tagrfp-caax* larvae, we observed that myelinated nerve volume was significantly reduced in 7 dpf *cd59^uva48^* mutant larvae (Figure 5 – figure supplement 1B, E). Because axon volume was unaltered, as indicated by acetylated α-Tubulin labeling (Figure 5 – figure supplement 1C, F), we hypothesized that myelination was affected in *cd59^uva48^* mutants. Utilizing TEM, we compared the myelin ultrastructure in pLLNs from wildtype and *cd59^uva48^* mutant larvae at 7 dpf. From these data, we observed a decrease in the number of myelin wraps around each axon in *cd59^uva48^* mutant larvae compared to wildtype siblings (Figure 5A, D). Because the total number of myelinated axons did not change between groups (Figure 5 – figure supplement 1G), these findings indicate that the reduction in myelin volume is due to, at least in part, fewer wraps of myelin around each axon. Overall, these data demonstrate that Cd59-dependent regulation of SC proliferation is necessary for myelin development.

**Figure 5.**
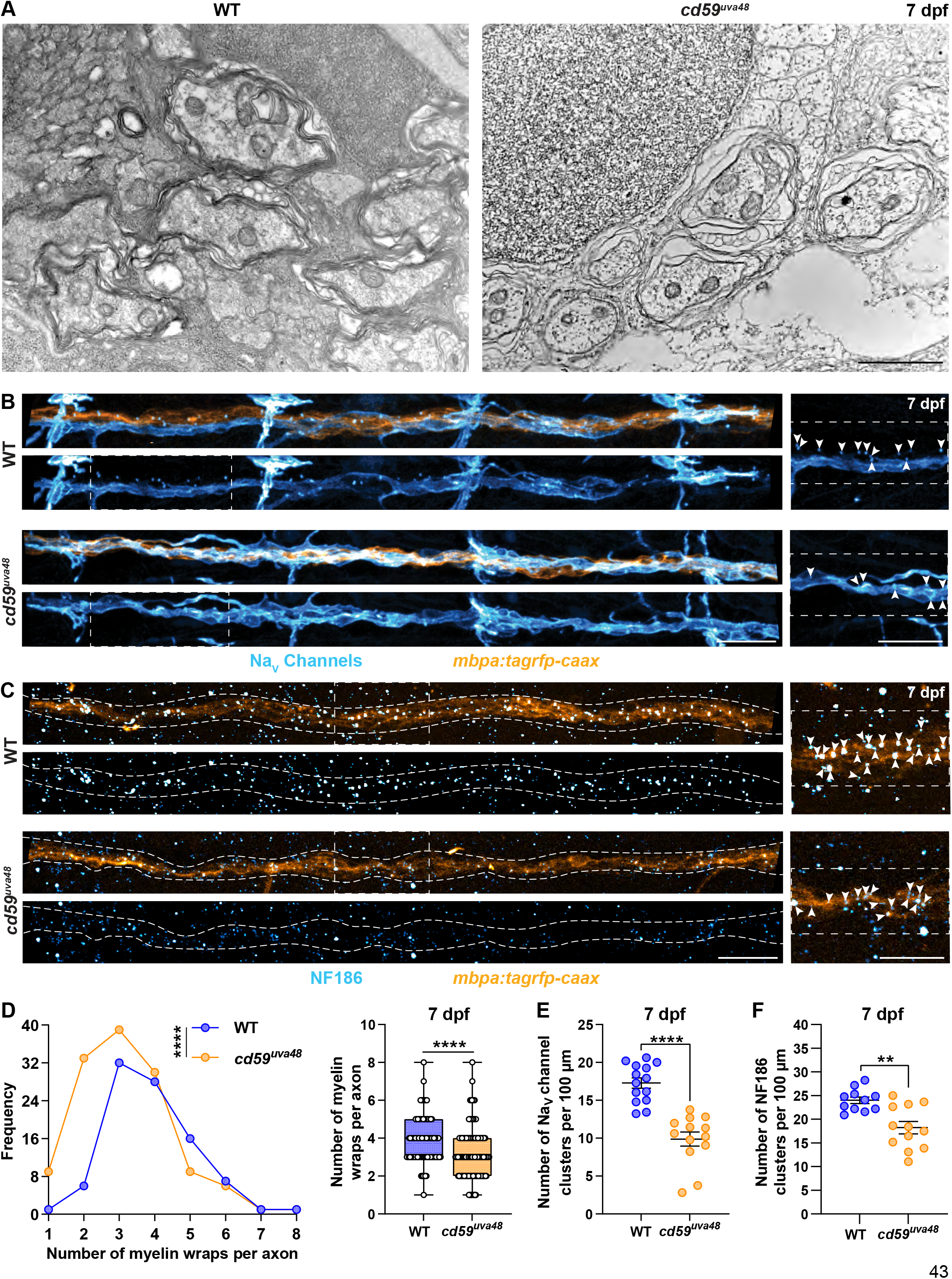
Myelin and node of Ranvier development is impaired in *cd59^uva48^* mutants. **(A)** Transmission electron micrographs showing pLLN axons myelinated by SCs at 7 dpf. **(B)** IF showing NaV channels (cyan) along *mbpa:tagrfp-caax*-positive pLLNs (orange) at 7 dpf. Diffuse NaV channel staining along unmyelinated nerves was not quantified. White dashes lines outline the pLLN. White dashed boxes correspond with the insets on the right. (**C**) IF showing NF186 clusters (cyan) along *mbpa:tagrfp-caax*-positive pLLNs (orange) at 7 dpf. White dashed lines outline the pLLN, and the white dashed boxes correspond to the insets on the right. Representative images in (**B**) and (**C**) depict somites 11 to 13 (∼320 µm). (**D**) Frequency distribution (right) and box and whisker plot (left) of the number of myelin wrappings per pLLN axon at 7 dpf. Data was collected from three sections per fish separated by 100 µm (mean ± SEM: WT: 3.89 ± 0.1, *cd59^uva48^*: 3.08 ± 0.2; p < 0.0001; dot = 1 myelinated axon; n = 3 fish per group). Data quantified in (**D**) were determined from electron micrographs in (**A**). (**E**) Scatter plot of the number of Na_V_ channel clusters along *mbpa:tagrfp*-positive pLLN nerves at 7 dpf (mean ± SEM: WT: 17.3 ± 0.7, *cd59^uva48^*: 9.9 ± 0.9; p < 0.0001; dot = 1 fish). (**F**) Scatter plot of the number of NF186 clusters along *mbpa:tagrfp*-positive pLLN nerves at 7 dpf (mean ± SEM: WT: 24.0 ± 0.7, *cd59^uva48^*: 18.2 ± 1.3; p = 0.0011; dot = 1 fish). Data were collected from somites 3 to 13 (∼320 µm) and normalized to units per 100 µm for (**E**) and (**F**). Scale bars: (**A**) 1 µm; (**B, C**) 25 µm. **<u>Supplemental and Source Data for Figure 5</u>** **Figure 5—Figure Supplement 1.** Myelin volume is reduced in *cd59^uva48^* mutants. **Figure 5 – source data 1.** Source data for quantification of myelin wraps. **Figure 5 – source data 2.** Source data for quantification of Na_V_ channel clusters. **Figure 5 – source data 3.** Source data for quantification of NF186 clusters. **Figure 5 – figure supplement 1 – source data 1.** Source data for quantification of myelin sheath length. **Figure 5 – figure supplement 1 – source data 2.** Source data for quantification of myelinated nerve volume. **Figure 5 – figure supplement 1 – source data 3.** Source data for quantification of axon volume. **Figure 5 – figure supplement 1 – source data 4.** Source data for quantification of myelinated axons.

In addition to forming myelin, myelinating SCs collaborate with axons to construct nodes of Ranvier, which occur between adjacent myelin sheaths and are essential for rapid neurotransmission along myelinated axons (Rasband and Peles, 2021). Therefore, we were curious if perturbed myelin formation in *cd59^uva48^* mutant larvae would also impact nodal development. An important aspect of node construction is the clustering of axonal sodium channels, which assists in the saltatory conduction of action potentials along the nerve and is facilitated by interactions between SC-associated gliomedin and neuronal cell adhesion molecule (NrCAM) and axonal neurofascin 186 (NF186) (Eshed et al., 2005; Feinberg et al., 2010; Rasband and Peles, 2021; Susuki et al., 2013). To investigate assembly of the node, we labeled *mbpa:tagrfp-caax* larvae with antibodies to visualize sodium channels and NF186 at 7 dpf. These studies revealed that *cd59^uva48^* mutant larvae had fewer sodium channel clusters along the pLLN (Figure 5B, E). Accordingly, mutant nerves also had fewer clusters of NF186 (Figure 5C, F). These findings indicate that NF186 fails to properly assemble at nodes of Ranvier in *cd59^uva48^* mutant larvae which in turn leads to impaired sodium channel clustering. Overall, this data shows that regulating SC proliferation through Cd59 is important for myelin formation and node of Ranvier assembly during development.

### Developmental inflammation stimulates Schwann cell proliferation and is regulated by Cd59

Cd59 is best known for its ability to inhibit complement-dependent cell lysis, protecting healthy cells from premature death during times of inflammation, such as during an infection or after an injury (Davies et al., 1989; Mead et al., 2004; Stahel et al., 2009; Yao and Verkman, 2017). Interestingly, at sublytic levels, complement can stimulate SC and OLC proliferation *in vitro* without inducing cell death (Dashiell et al., 2000; Hila et al., 2001; Rus et al., 1997, 1996; Tatomir et al., 2020). Complement is also a potent driver of inflammation, which is also known to drive cell proliferation (Kiraly et al., 2015; Larson et al., 2020; Morgan, 2016; Morgan and Harris, 2015; Silva et al., 2020). Although complement is present (Magdalon et al., 2020; Zhang and Cui, 2014), it is unclear if developmental levels of complement impact SC proliferation *in vivo* and whether this process is Cd59-dependent.

To determine if complement activity is increased in *cd59^uva48^* mutant larvae, we first looked at changes in membrane attack complex (MAC)-binding in developing SCs. MACs are comprised of complement proteins C5b, C6, C7, C8, and C9 and represent the culmination of the three complement pathways (classical, lectin, and alternative) (Bayly-Jones et al., 2017). During MAC formation, C9 will polymerize to form pores in the cell membrane, inducing cell proliferation or cell death depending on the concentration of pores (Bayly-Jones et al., 2017; Morgan, 1989; Tegla et al., 2011). In healthy cells, Cd59 will bind to C8 and C9 and prevent polymerization of C9 as well as subsequent pore formation (Meri et al., 1990; Ninomiya and Sims, 1992; Rollins et al., 1991). We hypothesized that if Cd59 was dysfunctional, we would see an increase in MAC formation in SC membranes. Using an antibody specific to C5b8-C5b9, which recognizes assembled MACs, we observed that *cd59*^uva48^ mutant larvae had more MACs localized to *sox10:megfp*-positive SC membranes compared to wildtype controls at 55 hpf (Figure 6A, B), demonstrating that developing SCs are no longer protected from complement in *cd59^uva48^* mutants.

**Figure 6.**
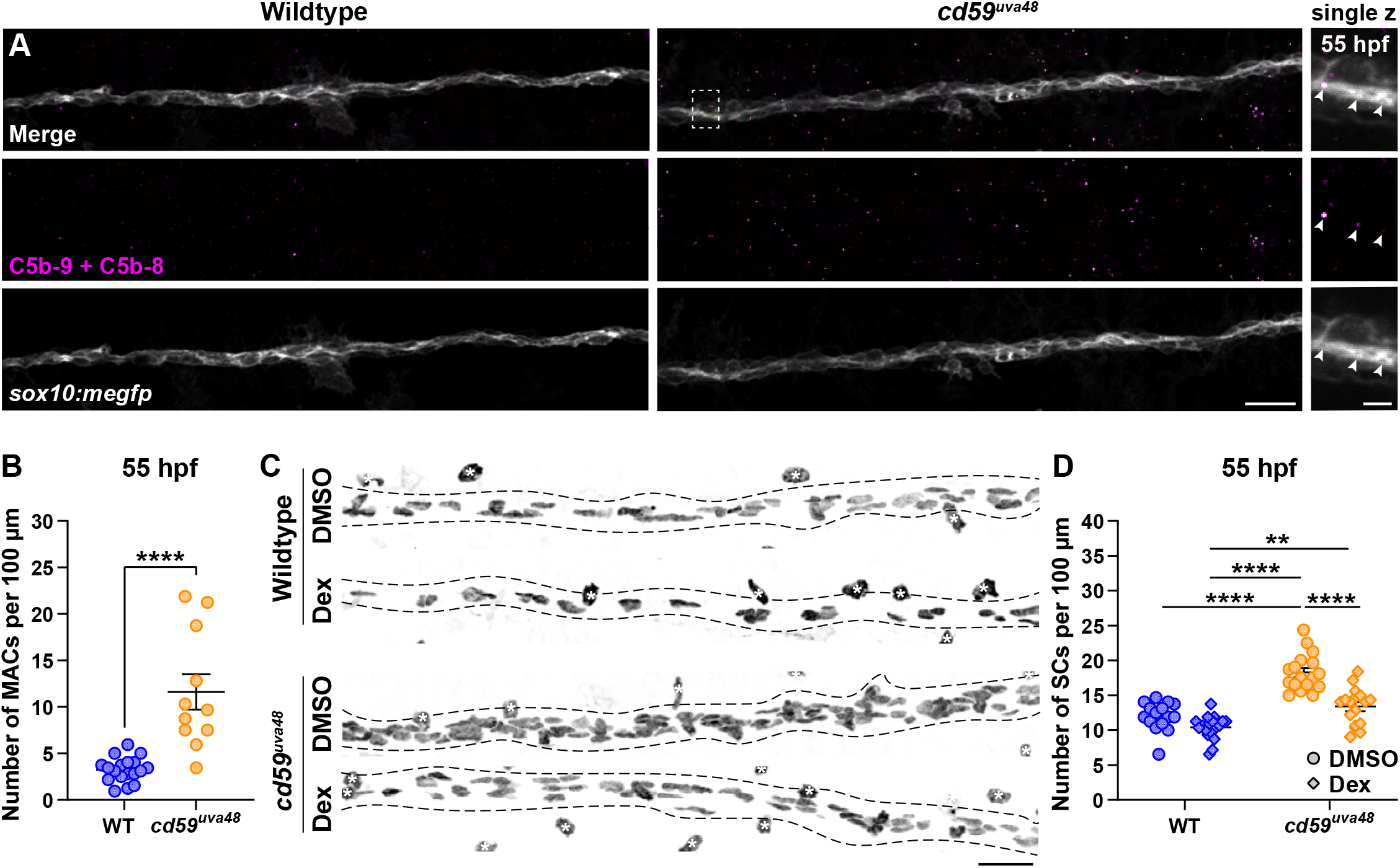
Cd59 and inflammation regulate developmental SC proliferation. (**A**) IF showing MACs (C5b-9+C5b-8; magenta) in *sox10:megfp-*positive pLLN SC membranes (grey) at 55 hpf. White dotted box corresponds with inset of a single z-plane on the right showing that MACs are within SC membranes. (**B**) Scatter plot of the number of MACs in SC membranes at 55 hpf (mean ± SEM: WT: 3.3 ± 0.3, *cd59^uva48^*: 11.6 ± 1.9; p < 0.0001; dot = 1 fish). (**C**) IF showing Sox10-positive pLLN SCs (black/grey) at 55 hpf in embryos treated with DMSO or 100 µM Dex. Black dashed lines outline the pLLN. Sox10-positive pigment cells outside of the dashed lines (white asterisks) were not included in the analysis. (**D**) Scatter plot of the number of pLLN SCs at 55 hpf (mean ± SEM: DMSO: WT: 12.0 ± 0.5, *cd59^uva48^*: 18.1 ± 0.6; Dex: WT: 10.4 ± 0.4, *cd59^uva48^*: 13.4 ± 0.6; p values: WT DMSO vs *cd59^uva48^* DMSO: p <0.0001, WT Dex vs *cd59^uva48^* DMSO: p < 0.0001, WT Dex vs *cd59^uva48^* Dex: p = 0.0014, *cd59^uva48^* DMSO vs *cd59^uva48^* Dex: p < 0.0001; dot = 1 fish). All data were normalized to units per 100 µm. Scale bars: (**A**) 10 µm; inset, 5 µm; (**C**) 25 µm. **<u>Supplemental and Source Data for Figure 6</u>** **Figure 6—Figure Supplement 1.** Dexamethasone does not interfere with development. **Figure 6 – source data 1.** Source data for quantification of SC-associated MACs. **Figure 6 – source data 2.** Source data for quantifications of SCs after dexamethasone treatment. **Figure 6—Figure Supplement 1. Dexamethasone does not interfere with development.** (**A**) Bright-field images of wildtype and mutant embryos (55 hpf) treated with DMSO or 100 µM Dex from 24 to 55 hpf. Scale bar: 0.25 mm.

To determine if increased complement activity and inflammation were responsible for excessive SC proliferation, we treated *cd59^uva48^* mutant and wildtype embryos with dexamethasone (Dex), a glucocorticoid steroid agonist known to inhibit inflammation and associated complement activity (Engelman et al., 1995; Silva et al., 2020). To do this, wildtype and mutant embryos were incubated in 1% DMSO or 1% DMSO plus 100 μM Dex from 24 to 55 hpf. This treatment method had no notable impacts on anatomical or behavioral development (Figure 6 – figure supplement 1A). After fixing the embryos, the number of pLLN SCs was quantified using a Sox10 antibody. Excitingly, Dex treatment in *cd59^uva48^* mutant embryos restored SC numbers to wildtype levels, whereas wildtype SCs were not significantly affected by Dex application (Figure 6C, D). Together, this data demonstrates that developmental inflammation aids in normal SC proliferation and that this process is amplified when *cd59* is mutated.

Previously, we showed that excessive SC proliferation interfered with myelin and node of Ranvier development (Figure 5 and Figure 5 – figure supplement 1). We therefore hypothesized that inhibiting inflammation-induced SC proliferation could rescue these aspects of nerve development. To investigate this hypothesis, we treated *cd59^uva48^* mutant embryos with 1% DMSO or 1% DMSO plus 100 μM Dex from 24 to 75 hpf to encompass most developmental SC proliferation. The larvae were then transferred to PTU-egg water and raised until 7 dpf. Labeling with a Sox10 antibody confirmed that this treatment regimen restored SC proliferation similarly to that described in Figure 6C and 6D (Figure 7 – figure supplement 1A, B). We then repeated the same treatment procedure in wildtype and *cd59^uva48^* mutant *mbpa:tagrfp-caax* embryos and quantified sodium channel antibody labeling at 7 dpf. Remarkably, we observed that dexamethasone treatment dramatically increased the number of sodium channel clusters in *cd59^uva48^* mutant larvae, achieving cluster levels similar to wildtype siblings (Figure 7A, C). Similarly, when comparing myelinated nerve volume in *mbpa:tagrfp-caax* larvae at 7 dpf, we saw that dexamethasone could also restore *cd59^uva48^* mutant nerve volume to wildtype type levels (Figure 7B, D). Collectively, these data demonstrate that inflammation-induced SC proliferation contributes to perturbed myelin and node of Ranvier development. Furthermore, inhibition of developmental inflammation can protect nerve development after *cd59* mutation.

**Figure 7.**
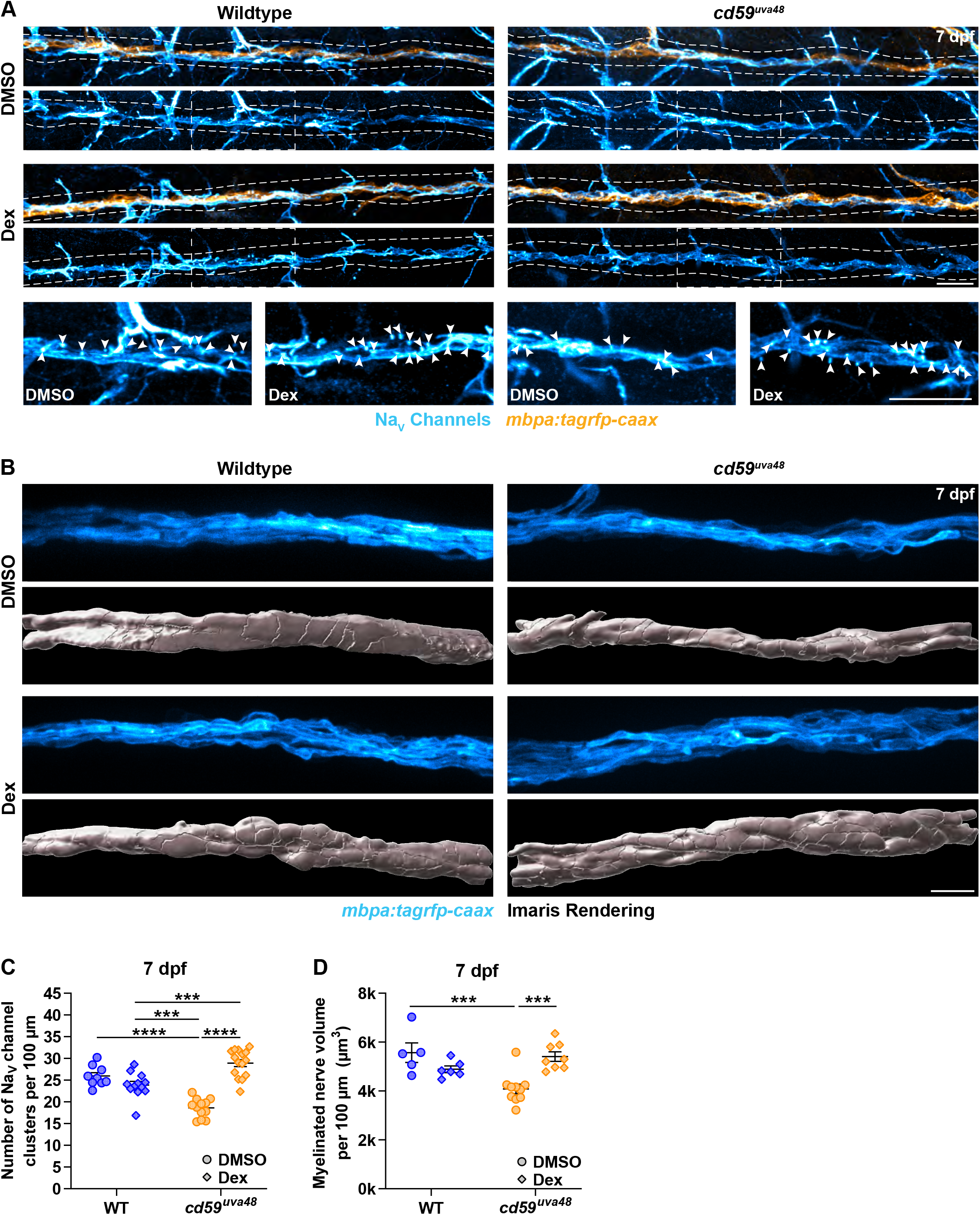
Cd59 and inflammation regulate myelin and node of Ranvier development. (**A**) IF showing Na_V_ channels (cyan) along *mbpa:tagrfp-caax*-positive nerves (orange) at 7 dpf in larvae treated with DMSO or 100 µM Dex. Diffuse Na_V_ channel staining along unmyelinated nerves was not quantified. White dashes lines outline the pLLN. White dashed boxes correspond with the insets below. Representative images are from somite 11 to 13 (∼320 µm). (**B**) *In vivo* imaging showing the volume of *mbpa:tagrfp-caax-*positive nerves at 7 dpf in larvae treated with DMSO or 100 µM Dex. Bottom panels depict Imaris renderings (white) of myelinated nerve volumes. Representative images are from somite 12 (∼110 µm). (**C**) Scatter plot of the number of Na_V_ channel clusters along *mbpa:tagrfp-caax*-positive nerves at 7 dpf (mean ± SEM: DMSO: WT: 26.0 ± 0.8, *cd59^uva48^*: 18.6 ± 0.6; Dex: WT: 23.9 ± 0.8, *cd59^uva48^*: 28.9 ± 0.8; p values: WT DMSO vs *cd59^uva48^* DMSO: p < 0.0001, WT Dex vs *cd59^uva48^* DMSO: p = 0.0001, WT Dex vs *cd59^uva48^* Dex: p = 0.0001, *cd59^uva48^* DMSO vs *cd59^uva48^* Dex: p < 0.0001; dot = 1 fish). Data were collected from somites 3 to 13 (∼320 µm). (**D**) Scatter plot of myelinated nerve volumes at 7 dpf (mean ± SEM: DMSO: WT: 5.6 ± 0.4, *cd59^uva48^*: 4.1 ± 0.2; Dex: WT: 4.9 ± 0.1, *cd59^uva48^*: 5.4 ± 0.2; p values: WT DMSO vs *cd59^uva48^* DMSO: p = 0.0009, *cd59^uva48^* DMSO vs *cd59^uva48^* Dex: p = 0.0006; dot = 1 fish). Data were collected from somite 12 (∼110 µm). All data were normalized to units per 100 µm. Scale bars: (**A**) 25 µm; inset, 25 µm; (**B**) 10 µm. **<u>Supplemental and Source Data for Figure 7</u>** **Figure 7—Figure Supplement 1.** Extended dexamethasone treatment had same impact on SC proliferation in *cd59^uva48^* mutant larvae. **Figure 7 – source data 1.** Source data for quantification of Na_V_ channel clusters after dexamethasone treatment. **Figure 7 – source data 2.** Source data for quantification of myelinated nerve volume after dexamethasone treatment. **Figure 7 – figure supplement 1 – source data 1.** Source data for quantification of SCs after extended dexamethasone treatment.

## Discussion

Myelination during nervous system development is essential for neural function, providing trophic and structural support to axons as well as quickening electrical conduction (Ritchie, 1982; Stadelmann et al., 2019). Consequently, impairment of this process can be devastating to patient quality of life (Stadelmann et al., 2019; van der Knaap and Bugiani, 2017). For this reason, the genetic mechanisms that orchestrate myelinating glial cell development requires continued exploration. Over the past few decades, we and others have noted expression of *cd59* that is conserved in developing myelinating glial cells across multiple organisms, including zebrafish, rodents, and humans (Gerber et al., 2021; Howard et al., 2021; Marisca et al., 2020; Marques et al., 2018, 2016; Piller et al., 2021; Saunders et al., 2019; Siems et al., 2021; Sun et al., 2013; Wolbert et al., 2020; Zajicek, 1995; Zhu et al., 2019). Despite these observations, there had been little exploration into the function of Cd59 in the developing nervous system. In this study, we demonstrate that *cd59* is expressed in a subset of developing SCs as well as mature OLs and SCs, revealing transcriptional heterogeneity among myelinating glial cells during development. Focusing on SCs, we demonstrated that Cd59 regulates SC proliferation induced by developmental inflammation and that this process is necessary to ensure myelin and node of Ranvier assembly during myelinated nerve formation. Overall, these findings illuminate the intersection of the innate immune system and glial cells and how they collaborate to establish a functioning nervous system during development.

Here, we showed that Cd59-limited proliferation is elicited by developmental inflammation. This finding provokes many interesting questions. First, these data reiterate the idea that the innate immune system and genes traditionally active in immune cells are first used during development to guide nervous system assembly and formation. In the CNS, there is evidence that complement aids in stimulating synaptic pruning of developing dendrites, directing cell polarity in the ventricular zone, guiding cortical neuron migration, and fostering neural progenitor cell proliferation (Coulthard et al., 2018, 2017; Denny et al., 2013; Gorelik et al., 2017; Magdalon et al., 2020; Paolicelli Rosa C. et al., 2011; Stevens et al., 2007). Similarly, inflammasome signaling was recently shown to be a necessary asset in Purkinje neuron development and mutations in this pathway are associated with increased DNA damage and behavioral deficits (Lammert et al., 2020). Finally, microglia, the resident innate immune cell of the CNS, have several roles in CNS development, including phagocytosing cell debris as well as pruning developing synapses and myelin (Hughes and Appel, 2020; Mazaheri et al., 2014; Silva et al., 2021; Stevens et al., 2007; Villani et al., 2019). To our knowledge, our study is the first examination of the role of inflammation and complement signaling in PNS development beyond the NCC stage (Carmona-Fontaine et al., 2011). These data prompt future exploration into this relationship between the nervous and immune systems during PNS formation.

During our investigation, we also show that *cd59* is expressed in a subset of SCs and OLs and is not expressed in other myelinating glial cells, including MEP glia and SCs that associate with spinal motor nerves. This expression pattern persists at least until 7 dpf in SCs, indicating that developmental heterogeneity lingers in the mature SCs. This finding provokes many topics for further investigation. First, how do *cd59*-positive SCs differ from *cd59*-negative SCs? Does this imply that a subset of SCs is more sensitive to complement activity, or are there other implications for these expression differences? Related, why do motor SCs lack *cd59*? Recent RNAseq analysis of developing satellite glial cells (SGCs) and SCs showed that glial precursors, SGCs, and iSCs are heterogenous and that this transcriptional diversity depended on the type of nerve/ganglia they associated with (Tasdemir-Yilmaz et al., 2021; Wiltbank and Kucenas, 2021). Motor neurons and sensory neurons are transcriptionally and functionally distinct and likely have different demands of the SCs that they are intimately associated with. Therefore, it follows that sensory SC functionality may require Cd59 whereas motor SCs do not. It will be interestingly to explore the consequences of this heterogeneity in future investigations.

Finally, we are also curious if Cd59 function is multi-faceted in the nervous system. Our findings demonstrated that Cd59 prevents overproliferation of SCs during development by shielding them from developmental inflammation. Similar observations have been noted in T cells and smooth muscle cells (Li et al., 2013; Longhi et al., 2005). That said, many other functions beyond proliferation control have been documented for CD59. For example, CD59 is required to instruct proximal-distal cell identity, a process that is necessary for proper cell positioning during limb regeneration (Echeverri and Tanaka, 2005). In this study, we noted that floor plate and hypochord cells express *cd59*. During neural tube development, floor plate cells play an important role in determining cell fate as well as dorsal-ventral patterning in the spinal cord (Hirano Shigeki et al., 1991; Yu et al., 2013), whereas the hypochord orchestrates midline blood vessel pattering (Cleaver and Krieg, 1998). Considering the role of CD59 in dictating proximal-distal cell identity during limb regeneration (Echeverri and Tanaka, 2005), it would be interesting to see if Cd59 participated in similar signaling pathways in floor plate and hypochord cells. CD59a also helps facilitate vesicle signaling, which seems to be particularly important during insulin release and is implicated in diabetic patients (Golec et al., 2019; Krus et al., 2014). Within the nervous system, vesicle-dependent signaling is important for myelin formation as well as trafficking within the myelin sheath (Baron and Hoekstra, 2010; Reiter and Bongarzone, 2020; White and Krämer-Albers, 2014). Electron micrographs show that CD59a is positioned throughout the myelin sheath in oligodendrocytes (Siems et al., 2021). Notably, CD59a molecules are observed deep within compacted myelin (Siems et al., 2021). Because Cd59 is unlikely to encounter complement when it is so isolated from the extracellular environment, we hypothesize that Cd59 is playing an additional role in mature myelinating glial cells, possibly by regulating vesicle signing within the myelin sheath. Ultimately, there remains many areas of exploration in fully characterize the role of Cd59 in the developing and mature nervous system.

## Materials and Methods

### Key Resources Table

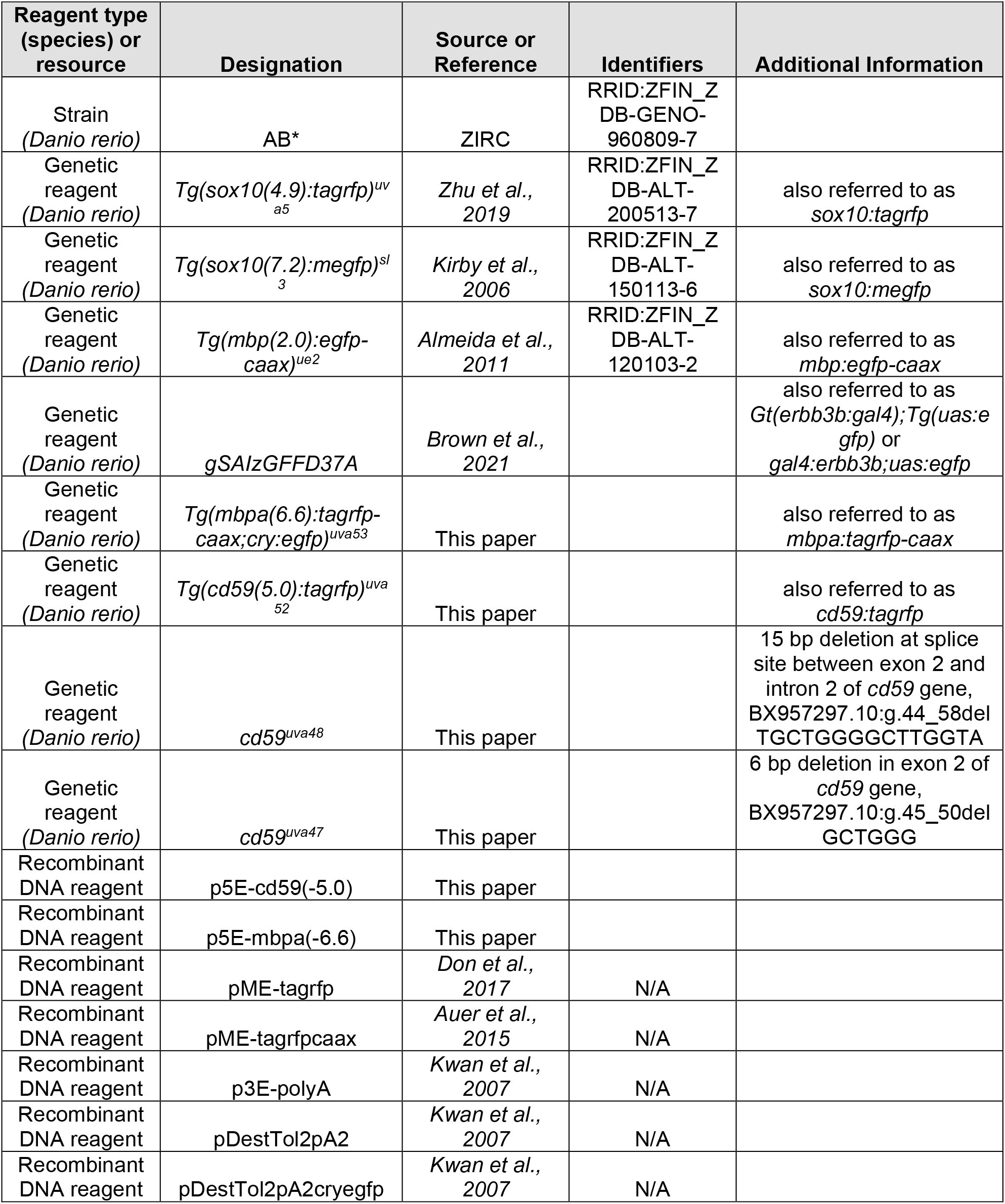
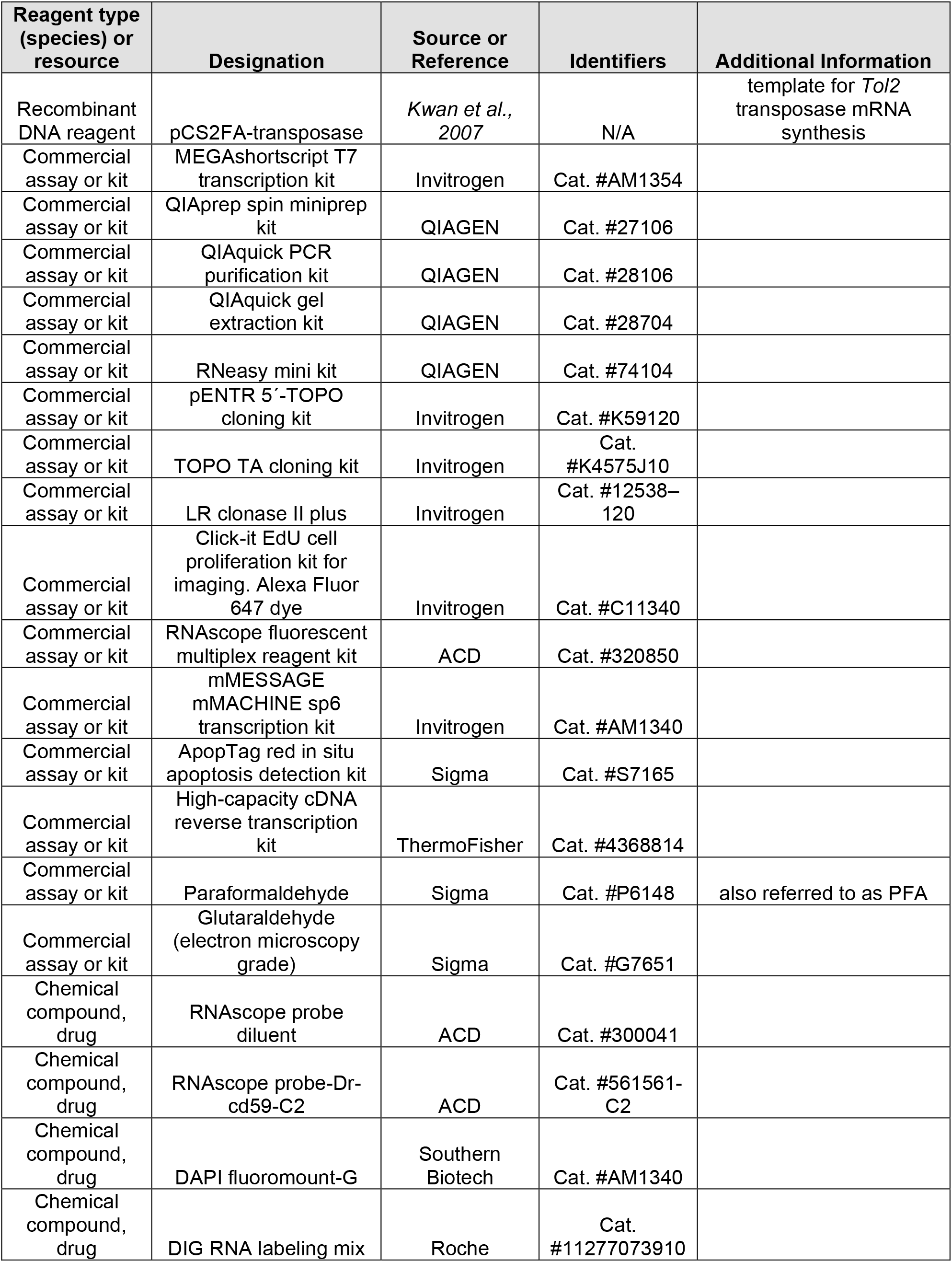
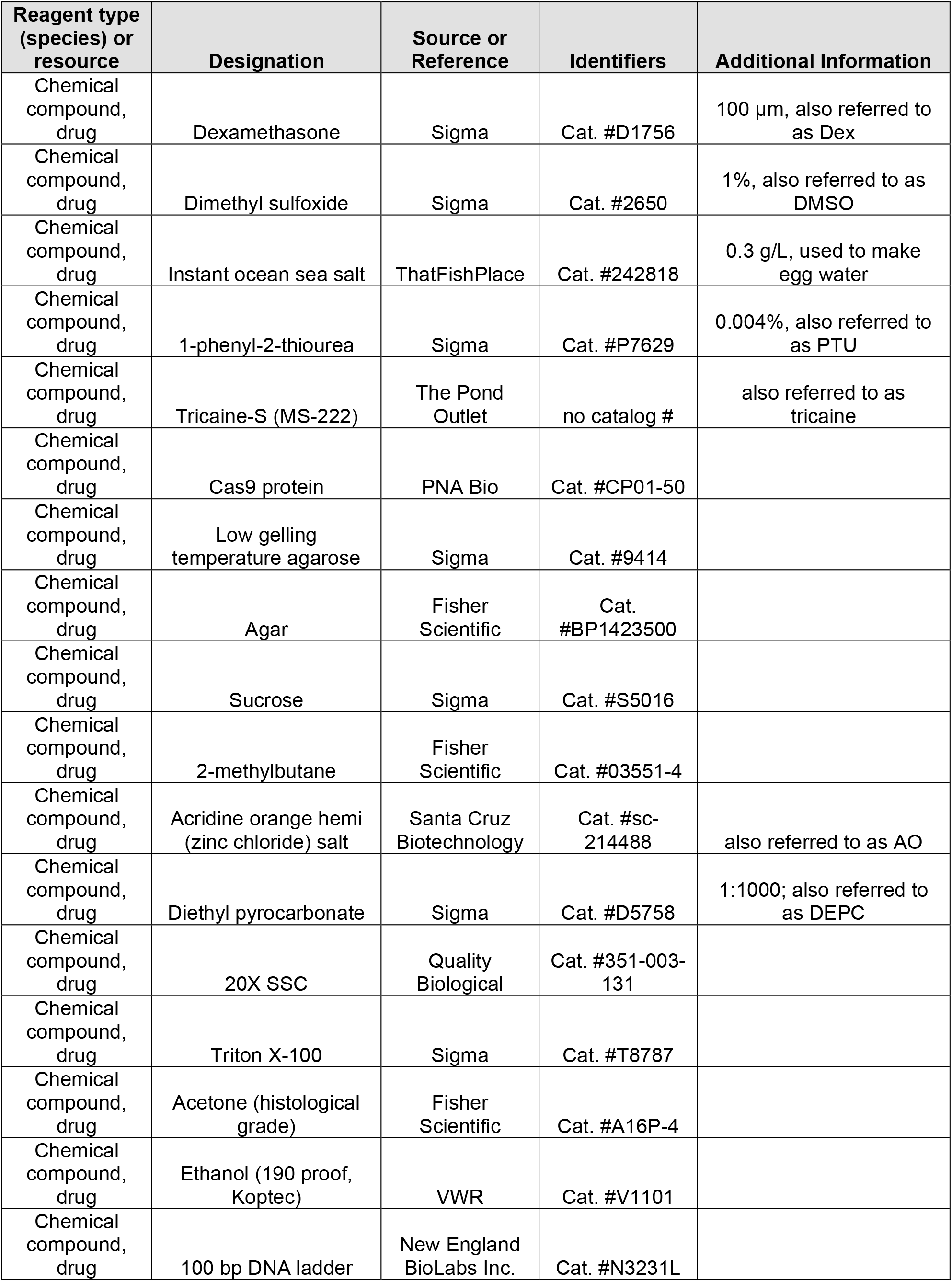
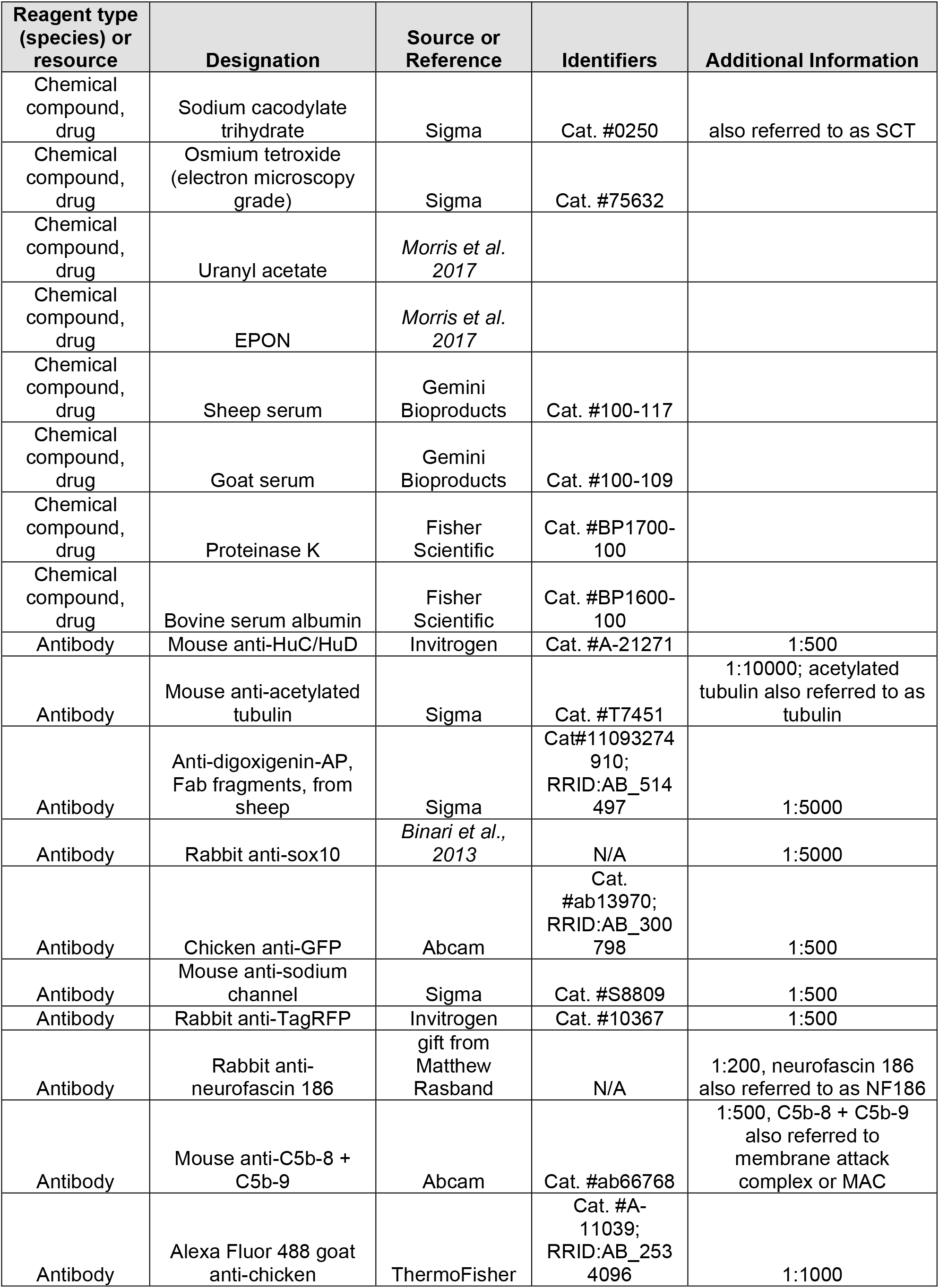
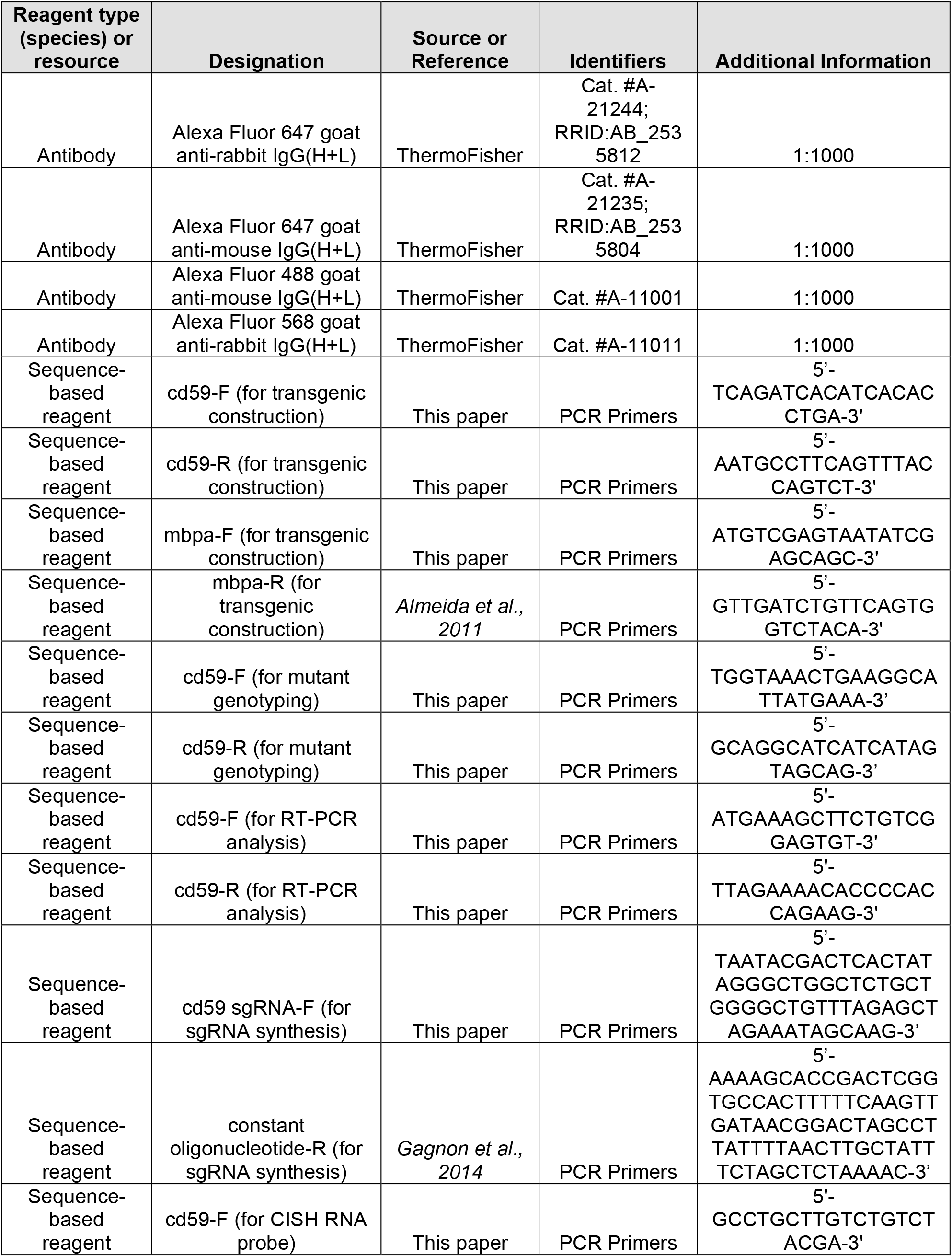
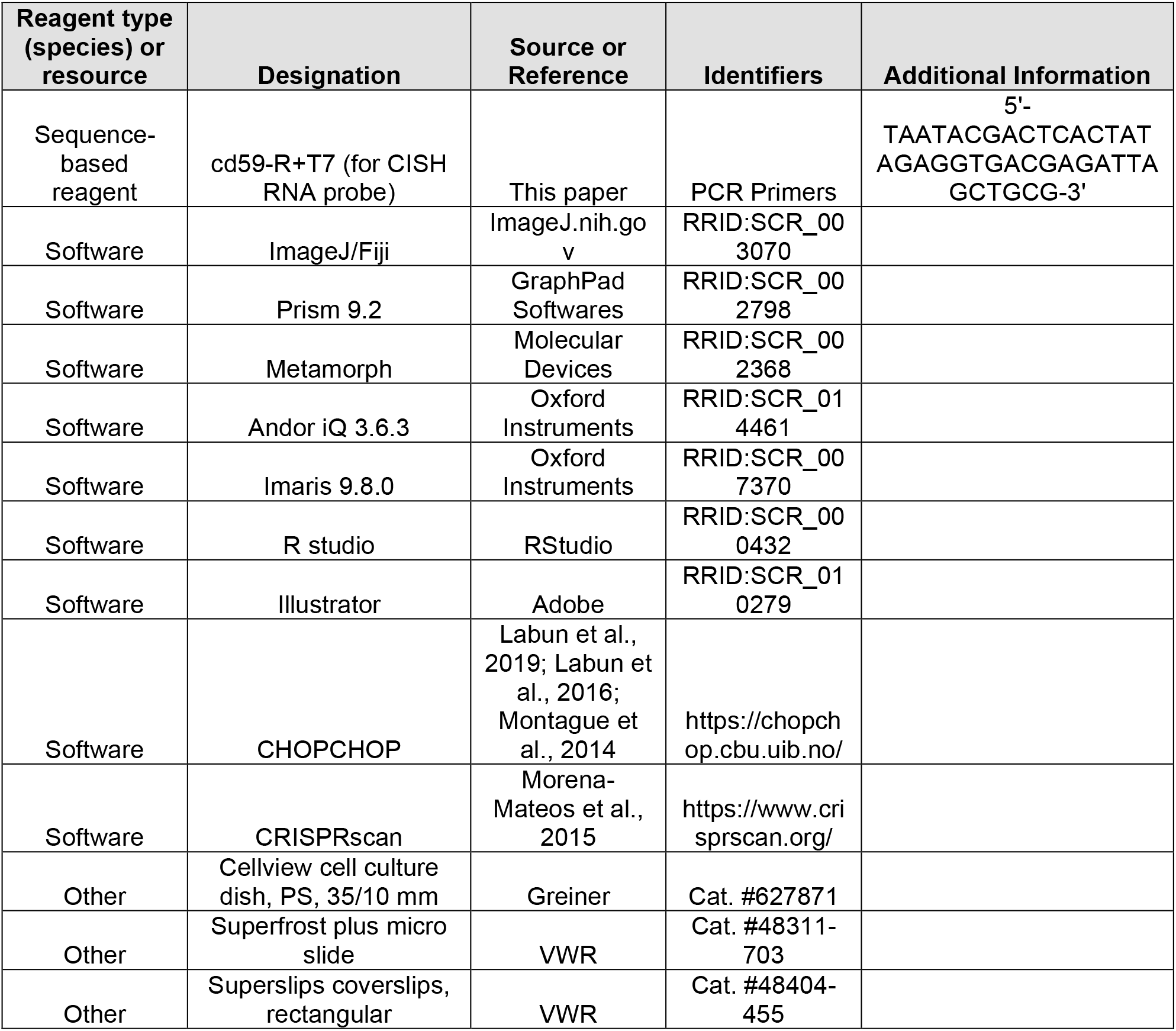

### Zebrafish Husbandry

All animal studies were approved by the University of Virginia Institutional Animal Care and Use Committee. Adult zebrafish were housed in tanks of 8 to 10 fish/L in 28.5 °C water. Pairwise mating of adult zebrafish generated zebrafish embryos for all experiments. The embryos were raised in egg water (0.3 g instant ocean sea salt per L reverse osmosis water) contained in 10 cm petri dishes and incubated at 28.5 °C. Embryos used for experiments were staged by hours post fertilization or days post fertilization (hpf & dpf) (Kimmel et al., 1995). To minimize visual obstruction by pigmentation, egg water was exchanged for 0.004% 1-phenyl-2-thiourea (PTU; Sigma) in egg water at 24 hpf. Tricaine-S (MS-222; The Pond Outlet) was utilized as an anesthetic for embryos and larvae used in live imaging and euthanasia. Embryo and larvae sex were undetermined for all experiments because sex cannot be ascertained until ∼25 dpf in zebrafish (Takahashi, 1977). To maintain genetic diversity, transgenic lines were renewed through outcrossing.

### Zebrafish Transgenic Lines

All transgene descriptions and abbreviations are included in the **Key Resources Table**. Transgenic lines produced during this study include: *Tg(cd59(5.0):tagrfp)^uva52^* and *Tg(mbpa(6.6):tagrfp-caax)^uva53^*. The methods used to generate these lines are described in **Generation of Transgenic Lines**. Previously published strains used in this study include: AB^*^, *Tg(sox10(7.2):megfp)^sl3^* (Kirby et al., 2006), *Tg(sox10(4.9):tagrfp)^uva5^* (Zhu et al., 2019), *Gt(erbb3b:gal4);Tg(uas:egfp)* (Brown et al., 2021) and *Tg(mbp(2.0):egfp-caax)^ue2^* (Almeida et al., 2011). All transgenic lines described are stable and incorporated into the germline. In addition to the stable lines, *cd59:tagrfp*, *mbpa:tagrfp-caax* and *mbp:egfp-caax* were also injected into embryos at the one-cell stage to create mosaic labeling (Kawakami, 2004).

### Zebrafish Mutant Lines

Mutant lines generated in the course of this study are as follows: *cd59^uva47^* (BX957297.10:g.45_50delGCTGGG) and *cd59^uva48^* (BX957297.10:g.44_58delTGCTGGGGCTTGGTA). The methods used to generate these lines are described in **Generation of Mutant Lines**. Mutant and wildtype fish were distinguished with PCR amplification and gel electrophoresis of the mutant allele (see **Generation of Mutant Lines**). Descriptions of all mutant lines and their abbreviations can be found the **Key Resources Table**.

### Generation of Transgenic Lines

The Tol2kit Gateway-based cloning system was used to produce transgenic constructs (Kwan et al., 2007). The following entry and destination vectors were used in the creation of these constructs: *p5E-cd59* (this paper)*, p5E-mbpa* (this paper), *pME-tagrfp-caax* (Auer et al., 2015), *pME-tagrfp* (Don et al., 2017), *p3E-polya* (Kwan et al., 2007), *pDestTol2pA2* (Kwan et al., 2007), and *pDestTol2pA2cryegfp*. These constructs were then used to produce the *Tg(cd59:tagrfp)* and *Tg(mbpa:tagrfp-caax)* zebrafish lines.

The *Tg(cd59(5.0):tagrfp)* line was generated as follows: PCR amplification of wildtype genomic DNA with forward primer, 5’-TCAGATCACATCACACCTGA-3’, and reverse primer, 5’-AATGCCTTCAGTTTACCAGTCT-3’, was used to clone 5 kb of the sequence upstream to the *cd59* gene (BX957297.10). The PCR product was purified through with the QIAquick gel extraction kit after gel electrophoresis (QIAGEN). The PCR product was subcloned into a pENTR 5’-TOPO vector (Invitrogen) to create the *p5E-cd59(−5.0)* vector. The resulting p5E-*cd59(5.0)* vector was transformed into chemically competent *E. coli* for amplification, isolated with the QIAprep spin miniprep kit (QIAGEN), and Sanger sequenced to verify accurate assembly. All Sanger sequencing described in the paper was conducted through GENEWIZ (Azenta Life Sciences; https://www.genewiz.com/en). LR reaction (Ashton et al., 2012) was used to ligate *p5E-cd59(5.0)* (this paper), *pME-tagrfp* (Don et al., 2017), *p3E-polya* (Kwan et al., 2007), and *pDestTol2pA2* (Kwan et al., 2007). The resulting *cd59(5.0):tagrfp* expression vector was transformed into chemically competent *E. coli* for amplification, isolated with the QIAprep spin miniprep kit (QIAGEN), and Sanger sequenced to verify accurate assembly. The stable *Tg(cd59(5.0):tagrfp)* line was established through co-microinjections of *cd59(5.0):tagrfp* expression vector (50 ng/µL) and *Tol2* transposase mRNA (20 ng/µL; mRNA synthesis described in **Generation of Synthetic mRNA**) in one-cell stage embryos (Kawakami, 2004). Founders were screened for germline incorporation of the transgene.

Similarly, the *Tg(mbpa(6.6):tagrfp-caax)* line was as described for the *Tg(cd59(5.0):tagrfp)* line except: PCR amplification of wildtype genomic DNA with forward primer, 5’-ATGTCGAGTAATATCGAGCAGC-3’ (this paper), and reverse primer, 5’- GTTGATCTGTTCAGTGGTCTACA-3’ (Almeida et al., 2011), was used to clone 6.59 kb of the sequence upstream to the *mbpa* gene (CU856623.7). The PCR product was subcloned into a pENTR 5’-TOPO vector (Invitrogen) to create the *p5E-mbpa(−6.6)* vector. LR reaction (Ashton et al., 2012) was used to ligate *p5E-mbpa(−6.6)* (this paper), *pME-tagrfp-caax* (Auer et al., 2015), *p3E-polya* (Kwan et al., 2007), and *pDestTol2pA2cryegfp* (Kwan et al., 2007). The final *mbpa(6.6):tagrfp-caax* construct and stable zebrafish lines were generated as described for *cd59(5.0):tagrfp*.

### Generation of Mutant Lines

*cd59* mutant lines generated during the course of this study were produced using CRISPR/Cas9 genome editing according to the methods described in Gagnon et al. 2014 except: The sgRNA targeting 5’-GTGCTGGCTCTGCTGGGGCTTGG-3’ in the second exon of *cd59* was identified with *CRISPRscan* (Morena-Mateos et al. 2015) and *Chop Chop* (Labun et al. 2019; Labun et al. 2016; Montague et al. 2014) and synthesized through PCR amplification with the following primers: forward primer, 5’-TAATACGACTCACTATAGGGCTGGCTCTGCTGGGGCTGTTTTAGAGCTAGAAATAG CAAG-3’, and constant oligonucleotide reverse primer, 5’- AAAAGCACCGACTCGGTGCCACTTTTTCAAGTTGATAACGGACTAGCCTTATTTTAA CTTGCTATTTCTAGCTCTAAAAC-3’ (Gagnon et al., 2014). All sgRNA synthesis and cleanup was performed as described in (Gagnon et al., 2014).

To generate a mutation in *cd59*, the sgRNA (200 ng/µL) was co-microinjected with Cas9 protein (600 ng/µL; PNA Bio) into embryos at the one-cell stage. Successful insertion and deletion mutation (INDEL) generation was verified according to the (Gagnon et al., 2014) protocol using the following primers: forward primer, 5’-TGGTAAACTGAAGGCATTATGAAA-3’, and reverse primer, 5’- GCAGGCATCATCATAGTAGCAG-3’.

To establish stable mutant lines, injected embryos were raised to adulthood and outcrossed with wildtype zebrafish. Founder offspring were screened for germline mutations through PCR amplification of *cd59* with the same genotyping primers listed above. To identify mutant alleles, the resulting PCR product was cloned into pCR4-TOPO TA vector (Invitrogen). Vectors containing *cd59* mutant alleles were amplified in chemically competent *E. coli*, isolated through miniprep (QIAGEN), and sequences were evaluated for INDELs. Founder offspring containing 6 bp (*cd59^uva47^*; BX957297.10:g.45_50delGCTGGG) and 15 bp deletions (*cd59^uva48^;* BX957297.10:g.44_58delTGCTGGGGCTTGGTA) in the *cd59* gene were selected for further experimentation and raised to establish stable mutant lines. All subsequent generations were genotyped as described in (Fontenas and Kucenas, 2021) with the same genotyping primers listed above, and the resulting PCR products were screen with gel electrophoresis on a 2.5% agarose gel.

### Generation of Synthetic mRNA

To aid in transgenic fish creation, *Tol2* transposase mRNA was transcribed from linearized pCS2FA-transposase (Kwan et al., 2007) with the mMESSAGE mMACHINE sp6 transcription kit (Invitrogen). The resulting mRNA was co-injected (20 ng/µL) with the transgenic constructs described in **Generation of Transgenic Lines**.

### RT-PCR Analysis

RNA was extracted with the RNeasy mini kit (QIAGEN) from 72 hpf *cd59^uva48^*, cd59^uva47^, and AB^*^ embryos. cDNA libraries were generated from the RNA using the high-capacity cDNA reverse transcription kit (ThermoFisher). Using the cDNA as a template, PCR amplification was performed with the following primers: forward primer, 5’- ATGAAAGCTTCTGTCGGAGTGT-3’,reverse primer, 5’- ATGAAAGCTTCTGTCGGAGTGT-3’. The resulting PCR products were visualized with gel electrophoresis on a 2.5% agarose gel.

To sequence the multiple mutant RNA transcripts found in the *cd59^uva48^* mutant, the PCR product was cloned into a pCR4-TOPO TA cloning vector (Invitrogen) as described in **Generation of Mutant Lines**. The isolated vectors were sequenced and analyzed for INDELs and premature termination codons (PTCs) with the Expasy protein translation tool (https://web.expasy.org/translate/)(Duvaud et al., 2021). *Cd59^uva48^* protein sequences were compared to sequences from *cd59^uva47^* and AB^*^ larvae.

### Confocal Imaging

All embryos were treated with egg water containing PTU (0.004%; Sigma) at 24 hpf to minimize pigmentation obstruction. For *in vivo* imaging, embryos and larvae (1 to 7 dpf) were dechorionated manually, if necessary, and anesthetized with 0.01% tricaine-S (MS-222) (The Pond Outlet). Low gelling temperature agarose (0.8%; Sigma) was used to immobilize the anesthetized fish in a 35 mm glass bottom dish (Greiner). Egg water containing PTU (0.004%; Sigma) and tricaine-S (MS-222) (0.01%; The Pond Outlet) was added to the dish prior to imaging to maintain anesthesia and suppress pigment production. For whole mount imaging of fixed fish (see **Immunofluorescence**), fixed embryos and larvae were immobilized with agarose in a glass bottom dish prior to imaging. For imaging tissue sections, sections were adhered to microscope slides prior to staining and imaging (see **Cryosectioning** and **Immunofluorescence**).

All fluorescent images were acquired with a 40x water immersion objective (NA = 1.1) mounted on a motorized Zeiss AxioObserver Z1 microscope equipped with a Quorum WaveFX-XI (Quorum Technologies) or Andor CSU-W (Andor Oxford Instruments plc.) spinning disc confocal system. Time-lapse experiments were imaged every 10 minutes for 7 to 24 hours, depending on the experiment. Z stacks were acquired at each time point for time-lapse imaging as well as single-time-point imaging for fixed whole mount and sectioned tissue (see **Immunofluorescence**, ***In Situ* Hybridization**, and **Cryosectioning**).

All experiments involving whole embryos and larvae were imaged with the 12^th^ somite at the center of the acquisition window to control for stage of anterior-posterior development. Exceptions include: (1) images of mosaic labeling with transgenic constructs, which were acquired regardless of anterior-posterior position, (2) images of Na_V_ channels and NF183, which were acquired from the 3^rd^ to the 13^th^ somite to ensure accurate assessment of nodes of Ranvier which are not evenly distributed along nerves, (3) images of the pLLG, which is anterior to the pLLN. All imaging of mosaic labeling as well as myelin volume quantification was obtained in live fish.

Images and videos were processed with either Metamorph (Molecular Devices) or IQ3 (Oxford Instruments). Fiji (ImageJ; imageJ.nih.gov), Imaris 9.8 (Oxford Instruments), and Illustrator (Adobe) were used for annotating videos and images, adjusting contrast and brightness, and data analysis.

### Cryosectioning

Fixed larvae were mounted in sectioning agar (1.5% agar; Fisher Scientific; 5% sucrose; Sigma; 100 mL ultrapure water) and cryopreserved in 30% sucrose in ultrapure water (Sigma) overnight at 4 °C. The agar blocks were frozen by placing them on a small raft floating on 2-methylbutane (Fisher Scientific), the container of which was submerged in a bath of liquid nitrogen. The blocks were sectioned to 20 µm with a cryostat microtome and mounted on microscope slides (VWR). The sections were stored at -20 °C until needed for immunofluorescence, *in situ* hybridization, or confocal imaging.

Prior to sectioning the offspring of heterozygous *cd59* mutant parents, the larvae were anesthetized in egg water with 0.01% tricaine-s (MS-222) (The Pond Outlet). The heads were removed with a razor blade. The heads were kept for genotyping, and the trunks were fixed for sectioning.

### Immunofluorescence

Larvae (24 hpf to 7 dpf) were fixed in 4% paraformaldyehyde (PFA; Sigma) for one hour shaking at room temperature (RT) for all experiments except Na_V_ channel and NF186 staining, in which samples were fixed for 30 minutes.

For whole mount imaging of embryo and larvae, the samples were prepared as described in (Fontenas and Kucenas, 2021). The following antibodies were used for whole mount immunofluorescence staining: mouse anti-acetylated tubulin (1:10000; Sigma), rabbit anti-sox10 (1:5000) (Binari et al., 2013), chicken anti-GFP (1:500; Abcam), mouse anti-sodium channel (1:500; Sigma), rabbit anti-TagRFP (1:500; Invitrogen), rabbit anti-neurofascin 186 (1:200; gift from Dr. Matthew Rasband), mouse anti-C5b-8 + C5b-9 (1:500; Abcam), Alexa Fluor 488 goat anti-chicken IgG(H+L) (1:1000; ThermoFisher), Alexa Fluor 647 goat anti-rabbit IgG(H+L) (1:1000; ThermoFisher), Alexa Fluor 647 goat anti-mouse IgG(H+L) (1:1000; ThermoFisher), Alexa Fluor 488 goat anti-mouse IgG(H+L) (1:1000; ThermoFisher), and Alexa Fluor 568 goat anti-rabbit IgG(H+L) (1:1000; ThermoFisher). Fish were immobilized and imaged in glass bottom dishes as described in **Confocal Imaging**.

For imaging of tissue sections, the samples were prepared as described in (Fontenas and Kucenas, 2021). The following antibodies were used for staining tissue sections: mouse anti-HuC/HuD (1:500; Invitrogen), rabbit anti-sox10 (1:5000)(Binari et al., 2013), Alexa Fluor 488 goat anti-mouse IgG(H+L) (1:1000; ThermoFisher), and Alexa Fluor 647 goat anti-rabbit IgG(H+L) (1:1000; ThermoFisher). After staining, all slides were mounted in DAPI fluoromount-G (Southern Biotech) and were coverslipped (VWR). The slides were stored in the dark until they were imaged as described in **Confocal Imaging**.

### *In Situ* Hybridization

#### Probe Synthesis

The *cd59* RNA probe for chromogenic *in situ* hybridization (CISH) was designed and synthesized in our lab. RNA was isolated from 3 dpf larvae with the RNeasy mini kit (QIAGEN). A cDNA library was generated from the 3 dpf RNA using the high-capacity cDNA reverse transcription kit (ThermoFisher). This cDNA was used as a template for the *cd59* RNA probe. The primers used to generate the *cd59* RNA probe were as follows: forward primer, 5’-GCCTGCTTGTCTGTCTACGA-3’, and reverse primer plus T7 sequence, 5’-TAATACGACTCACTATAGAGGTGACGAGATTAGCTGCG-3’. In addition to the *cd59* RNA probe, we also used previously published probes targeting *sox10* (Park and Appel, 2003) and *mbpa* (Brösamle and Halpern, 2002). CISH probe synthesis was performed as described in (Fontenas and Kucenas, 2021). For fluorescent *in situ* hybridization experiments (FISH), the *cd59* RNA probe (RNAscope probe-Dr-cd59-C2) was purchased from Advanced Cell Diagnostics (ACD).

#### Chromogenic In Situ Hybridization

Embryos and larvae (1 to 7 dpf) were dechorionated, if necessary, and fixed in 4% PFA for one hour shaking at RT and then transferred to 100% MeOH overnight at -20 °C. CISH was performed as described in (Hauptmann and Gerster, 2000). Images were obtained either using a Zeiss AxioObserver inverted microscope equipped with Zen, using a 40x oil immersion objective, or a Zeiss AxioObserver Z1 microscope equipped with a Quorum WaveFX-XI (Quorum Technologies) or Andor CSU-W (Andor Oxford Instruments plc.) spinning disc confocal system. Fiji (ImageJ; imageJ.nih.gov) and Illustrator (Adobe) were used for annotating images, adjusting contrast and brightness, and data analysis.

#### Fluorescent In Situ Hybridization

Larvae (1 to 7 dpf) were dechorionated, if necessary, and fixed with 4% PFA (Sigma) for one hour shaking at RT, dehydrated in 100% MeOH overnight at -20 °C, and cryosectioned (see **Cryosectioning**). The agar was gently removed by soaking the slides in 1X DEPC PBS [1:1000 diethyl pyrocarbonate (DEPC; Sigma) in 1X PBS. DEPC inactivated by autoclave].

To perform FISH, we used the RNAscope fluorescent multiplex reagent kit (ACD) with the following protocol modified from (Gross-Thebing et al., 2014): For all of subsequent steps, the slides kept in the dark and were coverslipped (VWR) during incubations except for during washes. To permeabilize the tissue, the sections were treated with Protease III (2 drops) and incubated at RT for 20 min. The tissue was gently rinsed three times with 1X DEPC PBS and washed with 1X DEPC PBS for 10 minutes at RT followed by an addition three rinses in 1X DEPC PBS. The sections were then hybridized with *cd59* probe (1:100 RNAscope probe-Dr-cd59-C2 in RNAscope probe diluent; ACD) in a 40 °C water bath overnight. The tissue was rinsed three times with 1X DEPC PBS and washed in 0.2X SSCT [0.2X SSC (Quality Biological) and 0.1% Triton X-100 (Sigma) in DEPC water (1:1000 DEPC in ultrapure water; DEPC inactivated by autoclave)] for 10 min and then rinsed again three times with 1X DEPC PBS. This rinse/wash routine was repeated between the following fixation and amplification steps: 4% PFA (5 min at RT), Amp1 (2 drops, 30 min at 40 °C), Amp2 (2 drops, 15 min at 40 °C), Amp3 (2 drops, 30 min at 40 °C), Amp4C (2 drops, 15 min at 40 °C), and DAPI (2 drops, 30 min at RT). After a final rinse/wash, the slides were coverslipped (VWR) with 0.2X SSCT and stored at 4 °C until imaged. Sections were imaged as described in **Confocal Imaging**.

### Cell Death Assays

#### Acridine Orange Incorporation Assay

To label cell death, 48 hpf embryos were labeled with acridine orange (AO) according to the protocol in (Lyons et al., 2005). Fish were immobilized and imaged in glass bottom dishes as described in **Confocal Imaging**.

#### TUNEL Assay

To label cell death, 48 hpf embryos were fixed in 4% PFA for one hour shaking at RT and then sectioned (see **Cryosectioning**). TUNEL staining was performed with the ApopTag red in situ apoptosis detection kit (Sigma) and followed by immunofluorescence for Sox10 (see **Immunofluorescence**). Sections were imaged as described in **Confocal Imaging**.

### EdU Incorporation Assay

To label mitotically-active cells, embryos were incubated in EdU [0.1 mM EdU (Invitrogen), 1% DMSO (Sigma), and egg water containing 0.004% PTU (Sigma)] from 48 to 55 hpf. The embryos were then fixed in 4% PFA for one hour shaking at RT and permeabilized as described in **Immunofluorescence**. We stained for EdU with the Click- it EdU cell proliferation kit for imaging (Alexa Fluor 647 dye; Invitrogen) as described in the kit protocol. The Click-it reaction was performed for one hour shaking at RT. The embryos were washed overnight shaking at 4 °C in 1X PBS. Fish were immobilized and imaged in glass bottom dishes as described in **Confocal Imaging**.

### Dexamethasone Treatment

To inhibit inflammation, embryos were incubated in 1% DMSO (Sigma) or 1% DMSO plus 100 µM dexamethasone (Dex; Sigma) at 28.5 °C. Dex concentration was chosen based on a previously published dose-response study in zebrafish larvae (Wilson et al., 2016, 2013). For embryos fixed at 55 hpf, the embryos were treated with DMSO or Dex from 24 to 55 hpf. For larvae fixed at 7 dpf, the embryos were treated from 24 to 75 hpf and then transferred to egg water containing PTU (0.004%; Sigma).

### Transmission Electron Microscopy

Headless larvae (7 dpf) were fixed for three days at 4 °C in EM fixation buffer [2% EM-grade glutaraldehyde (Sigma), 2% PFA (Sigma), 0.1M sodium cacodylate trihydrate, pH 7.3 (SCT; Sigma) in ultrapure water] and washed three times for 10 minutes each in 0.1M SCT. The samples were post-fixed in 1% osmium tetroxide (Sigma) in 0.1M SCT for one hour at 4 °C and washed three times for 5 min each in ultrapure water at RT. Contrast was initiated with 2% uranyl acetate for one hour at RT and washed four times for 5 min each in ultrapure water at RT. The samples were dehydrated as follows: 40% EtOH (2x 10 min), 60% (2x 10 min), 80% (2x 10 min), 100% (2x 10 min), acetone (2 min), and acetone/EPON (1:1, 15 min). The samples were then incubated in EPON at 4 C at 4 °C overnight with the lid open to allow for gas to escape. Samples were mounted in EPON and polymerized at 60 °C for 48 hours. Tissue blocks were sectioned transversely, stained, and imaged according to the methods described in (Morris et al., 2017). Sections were collected from three parts of the larvae, separated by 100 µm^*^, to enable quantification of three groups of SCs per pLLN.

### RNAseq Analysis

Published bulk and scRNAseq datasets from zebrafish and rodents were evaluated for *cd59* expression. The chosen datasets included analysis of some or many stages of myelinating glial cell development. Visualization and quantification of *cd59* expression was obtained through the applications included with the publications when possible (publications cited in Figure 1—Figure Supplement 1) with the following exceptions: (1) TPM quantification of bulk RNAseq of zebrafish myelinating glial cells was acquired from data analyzed in (Piller et al., 2021; Zhu et al., 2019). (2) Analysis of NCC lineage scRNAseq in zebrafish from the dataset presented in (Saunders et al., 2019) was performed as follows: Tissue collection, FACS, RT-PCR, single cell collection, library construction, sequencing, read alignment to Ensembl GRCz11, and preliminary data processing was performed as described by (Saunders et al., 2019), and the data was accessed through GEO via accession <u>GSE131136</u>. Dimensionality reduction and projection of cells in two dimensions was performed using uniform manifold approximation and projection (UMAP) (McInnes et al., 2018) and Louvain clustering (Blondel et al., 2008). Cell cluster visualization and analysis was performed using the FindMarkers, FeaturePlot, DimPlot, and VlnPlot functions of Seurat package 4.0.0 (Hao et al., 2021).

### Quantification and Statistical Analysis

#### Nerve Volume Quantification

Myelinated nerve and axon volume were quantified by creating a surface rendering of the imaged nerve using Imaris 9.8 (Oxford Instruments). The total volume of each surface rendering was quantified with Imaris 9.8 (Oxford Instruments).

#### Myelin Length Quantification

Myelin length was measured from images of mosaic *mbpa:tagrfp-caax* labeling using Fiji (ImageJ; imageJ.nih.gov). Specifically, the length of the myelin sheath was traced with the “freehand line” tool. The length was then quantified with the “measure” tool.

#### Axon Myelination Quantification

The number of myelin wraps around an axon as well as the number of myelinated axons per pLLN were measured in electron micrographs of pLLNs using Fiji (ImageJ; imageJ.nih.gov). Counting was performed with the “multi-point” and “measure” tools.

#### Na_V_ Channel, NF186, cd59 RNAscope, and MAC Quantification

Puncta quantification (specifically for quantification of Na_V_ channels, NF186, *cd59* RNAscope, and MAC quantification) was performed either in Fiji (ImageJ; imageJ.nih.gov) or Imaris (Oxford Instruments). To ensure only puncta within the pLLN were quantified, each z plane was quantified manually.

#### Cell Number Quantification

SC, neuron, and NCC images on the pLLN and in the CNS were quantified in Fiji (ImageJ; imageJ.nih.gov) by creating z projections and counting the number of cells with the “multi-point” and “measure” tools. pLLG neurons and NCCs were counted manually in Imaris 9.8 (Oxford Instruments).

#### Mitotic Event Quantification

Time-lapse images of developing SCs were evaluated for mitotic events (cell divisions) of *sox10:tagrfp*-positive SCs along the pLLN using Fiji (ImageJ; imageJ.nih.gov). Mitotic events were defined as follows: SC rounds up, chromatic condenses (darkening of the SC), and division of the daughter cells. The number of mitotic events were quantified with the “multi-point” and “measure” tools. Mitotic events in the representative videos were annotated arrows using the “draw arrow in movies” plugin (Daetwyler et al., 2020).

#### Statistical Analysis

Student t tests as well as one-way and two-way ANOVAs followed by Tukey’s multiple comparison tests, were performed using Prism 9.2 (GraphPad Softwares). The data in plots and the text are presented as means ± SEM.

#### Data Blinding

For offspring of heterozygous parents, embryos chosen for experimentation were blindly selected (wildtype and mutant embryos are indistinguishable by eye). Embryo genotype was revealed after imaging and before quantification. For offspring of homozygous parents, embryo genotype was known throughout the entire experiment.

### Artwork

All artwork was created in Illustrator (Adobe) by Ashtyn T. Wiltbank. SC and OL development figures (Figure 1A, B) were based on previous schematics and electron micrographs shown in (Ackerman and Monk, 2016; Cunningham and Monk, 2018; Jessen and Mirsky, 2005).

## Acknowledgments

We thank Lori Tocke for zebrafish care and members of the Kucenas Lab for valuable discussions. We thank Drs. Laura Fontenas, Xiaowei Lu, Sarah E. Siegrist, John R. Lukens, and Scott Zeitlin for helpful suggestions and training. We thank Drs. Laura Fontenas, Alev Erisir, Stacey J. Criswell, Natalia Dworak, and the Advanced Microscopy Facility for assistance with EM sample preparation and microscope training. We thank Dr. Matthew N. Rasband for the NF186 antibody. We thank Dr. David M. Parichy for his advice and for sharing scRNAseq data from (Saunders et al., 2019). This work was funded by NIH/National Institutes of Neurological Disorders and Stroke (NINDS): R01NS107525 (SK) and The Owens Family Foundation (SK).

## Additional Information

This work was funded by NIH/National Institutes of Neurological Disorders and Stroke (NINDS): R01NS107525 (SK) and The Owens Family Foundation (SK). The funders had no role in study design, data collection and interpretation, or the decision to submit the work for publication.

## Author Contributions

**Ashtyn T. Wiltbank:** Conceptualization, resources, data curation, formal analysis, validation, investigation, visualization, methodology, writing (original draft, review, and editing), and project administration

**Emma Steinson:** Investigation, validation, formal analysis

**Stacey J. Criswell:** Methodology

**Melanie Piller:** Data curation, formal analysis, visualization, writing (methodology draft)

**Sarah Kucenas:** Conceptualization, supervision, funding acquisition, validation, writing (original draft, review, and editing), and project administration

## Ethics

Animal experimentation: All animal studies were approved by the University of Virginia Institutional Animal Care and Use Committee, Protocol #3782.

## Figure Legends for Source Data

**Figure 1 – source data 1. Source data for *cd59* bulk, RNAseq expression.** Source data for *cd59* bulk, RNAseq expression depicted in Figure 1C. Data contributed to scatter plot of *cd59* expression (TPM) in OLCs, SCs, and neurons (N) at 72 hpf (left; mean ± SEM: OLC: 2145.1 ± 1215.1, SC: 40.1 ± 16.3; N: 240.5 ± 173.3; data point = replicate) as well as SCs at 36 and 72 hpf (right; mean ± SEM: 36 hpf: 0.0 ± 0.0, 72 hpf: 40.2 ± 16.3; data point = replicate).

**Figure 2 – source data 1. Source data for quantification of *cd59:tagrfp*-positive cells.** Source data for percent *cd59:tagrfp*-positive cells per 100 µm depicted in Figure 2E. Data contributed to scatter plot of percent *cd59:tagrfp*-positive SCs on the pLLN from 3 to 7 dpf (mean ± SEM: 3 dpf: 9.4 ± 1.5, 4 dpf: 10.0 ± 1.0, 5 dpf: 7.8 ± 1.2, 6 dpf: 4.4 ± 0.6, 7 dpf: 5.8 ± 0.6; p values: 3 dpf vs 6 dpf: p = 0.0126, 4 dpf vs 6 dpf: p = 0.0095, 4 dpf vs 7 dpf: p = 0.0477; data point = 1 fish). Data collected from somites 11 to 13 (∼320 μm) and normalized to units per 100 μm.

**Figure 3 – source data 1. Source data from quantification of *cd59* puncta.** Source data for the number of cd59 puncta in SCs at 72 hpf depicted in Figure 3H. Data contributed to scatter plot of the number of *cd59* RNA puncta in pLLN SCs (mean ± SEM: WT: 64.7 ± 7.6, *cd59^uva48^*: 14 ± 1.7; p < 0.0001; data point = 1 cell; n = 7 fish per group).

**Figure 3 – figure supplement 1 – source data 1. Source data from gel electrophoresis of RT-PCR of *cd59^uva48^* mutant embryos.** Unlabeled and labeled images of gel electrophoresis showing wildtype (357 bp) and *cd59^uva48^* (variable transcript size) RT-PCR products at 72 hpf. RT-PCR products were compared to 100 bp DNA.

**Figure 3 – figure supplement 1 – source data 2. Source data from gel electrophoresis of RT-PCR of *cd59^uva47^* mutant embryos.** Unlabeled and labeled images of gel electrophoresis showing wildtype (357 bp) and *cd59^uva47^* (351 bp) RT-PCR products at 72 hpf. RT-PCR products were compared to 100 bp DNA.

**Figure 4 – source data 1. Source data from quantification of pLLN SCs at 36 hpf.** Source data for the number of SCs on the pLLN at 36 hpf. Data contributed to scatter plot of the number of Sox10-positive SCs along the pLLN at 36 hpf (mean ± SEM: WT: 5.6 ± 0.2, *cd59^uva48^*: 9.1 ± 0.3; p value: p < 0.0001; data point = 1 fish). Data were collected from somites 11 to 13 (∼320 µm) and normalized to units per 100 µm.

**Figure 4 – source data 2. Source data from quantification of pLLN SCs at 48 hpf.** Source data for the number of SCs on the pLLN at 48 hpf. Data contributed to scatter plot of the number of Sox10-positive SCs along the pLLN at 48 hpf (mean ± SEM: WT: 8.6 ± 0.5, *cd59^uva48^*: 15.0 ± 0.7; p value: p < 0.0001; data point = 1 fish). Data were collected from somites 11 to 13 (∼320 µm) and normalized to units per 100 µm.

**Figure 4 – source data 3. Source data from quantification of pLLN SCs at 72 hpf.** Source data for the number of SCs on the pLLN at 72 hpf. Data contributed to scatter plot of the number of Sox10-positive SCs along the pLLN at 72 hpf (mean ± SEM: WT: 18.0 ± 0.4, *cd59^uva48^*: 24.3 ± 0.8; p value: p < 0.0001; data point = 1 fish). Data were collected from somites 11 to 13 (∼320 µm) and normalized to units per 100 µm.

**Figure 4 – source data 4. Source data from quantification of pLLN SCs at 7 dpf.** Source data for the number of SCs on the pLLN at 7 dpf. Data contributed to scatter plot of the number of Sox10-positive SCs along the pLLN at 7 dpf (mean ± SEM: WT: 27.0 ± 0.6, *cd59^uva48^*: 33.9 ± 0.8; p value: p < 0.0001; data point = 1 fish). Data were collected from somites 11 to 13 (∼320 µm) and normalized to units per 100 µm.

**Figure 4 – source data 5. Source data from quantification of EdU-positive SCs from 48 to 55 hpf.** Source data for the number of EdU-positive SCs on the pLLN from 48 to 55 hpf depicted in Figure 4G. Data contributed to scatter plot of the number of EdU-positive SCs along the pLLN at 55 hpf (mean ± SEM: WT: 6.7 ± 0.5, *cd59^uva48^*: 13.6 ± 0.5; p < 0.0001; dot = 1 fish). Data were collected from somites 11 to 13 (∼320 µm) and normalized to units per 100 µm.

**Figure 4 – source data 6. Source data for quantification of mitotic events from 48 to 55 hpf.** Source data for the number of mitotic events from 48 to 55 hpf depicted in Figure 4H. Data contributed to scatter plot of the number of mitotic events observed in SCs from 48 to 55 hpf (mean ± SEM: WT: 0.6 ± 0.1, *cd59^uva48^*: 1.7 ± 0.3; p = 0.0019; data point = 1 fish). Data were collected from somites 11 to 13 (∼320 µm) and normalized to units per 100 µm.

**Figure 4 – source data 7. Source data for quantification of mitotic events from 54 to 75 hpf.** Source data for the number of mitotic events from 48 to 55 hpf depicted in Figure 4I. Data contributed to scatter plot of the number of mitotic events observed in SCs from 54 to 75 hpf (mean ± SEM: WT: 4.1 ± 0.3, *cd59^uva48^*: 4.1 ± 0.4; data point = 1 fish). Data were collected from somites 11 to 13 (∼320 µm) and normalized to units per 100 µm.

**Figure 4 – source data 8. Source data for time of final division.** Source data for the time of final SC division during 54 to 75 hpf depicted in Figure 4J. Data contributed to scatter plot of the time of final cell division (hpf) observed in SCs from 54 to 75 hpf (mean ± SEM: WT: 69.1 ± 1.1, *cd59^uva48^*: 68.4 ± 0.9; data point = 1 fish). Data were collected from somites 11 to 13 (∼320 µm).

**Figure 4 – figure supplement 1 – source data 1. Source data for quantification of SCs in *cd59^uva47^* mutant embryos.** Source data for the number of SCs on the pLLN at 48 hpf in *cd59^uva47^* mutant embryos. Data contributed to scatter plot of the number of Sox10-positive SCs along the pLLN at 48 hpf (mean ± SEM: WT: 5.7 ± 0.3, *cd59^uva48^*: 8.3 ± 0.5; p = 0.0003; data point = 1 fish). Data was collected from somites 11 to 13 (∼320 µm) and normalized to units per 100 µm.

**Figure 4 – figure supplement 1 – source data 2. Source data for quantification of pLLG neurons.** Source data for the number of neurons per pLLG at 24 hpf depicted in Figure 4 – figure supplement 1E. Data contributed to scatter plot of the number of HuC/HuD-positive neurons in the pLLG at 24 hpf (mean ± SEM: WT: 20.9 ± 1.0, *cd59^uva48^*: 20.3 ± 1.1; data point = 1 fish).

**Figure 4 – figure supplement 1 – source data 3. Source data for quantification of pLLG-associated NCCs.** Source data for the number of NCCs associated with the pLLG at 24 hpf depicted in Figure 4 – figure supplement 1F. Data contributed to scatter plot of the number of Sox10-positive NCCs in the pLLG at 24 hpf (mean ± SEM: WT: 49.2 ± 2.1, *cd59^uva48^*: 50.8 ± 1.4; data point = 1 fish).

**Figure 4 – figure supplement 1 – source data 4. Source data for quantification of migrating NCCs.** Source data for the number of migrating NCCs per stream at 24 hpf depicted in Figure 4 – figure supplement 1H. Data contributed to scatter plot of the number of Sox10-positive, migrating NCCs in the trunk at 24 hpf (mean ± SEM: WT: 7.1 ± 0.4, *cd59^uva48^*: 7.3 ± 0.4; data point = 1 fish).

**Figure 4 – figure supplement 1 – source data 5. Source data for quantification of trunk neuromasts.** Source data for the number of trunk neuromasts at 7 dpf depicted in Figure 4 – figure supplement 1I. Data contributed to scatter plot of the number of Tubulin- positive neuromasts in the trunk at 7 dpf (mean ± SEM: WT: 7.4 ± 0.3, *cd59^uva48^*: 6.8 ± 0.3; data point = 1 fish).

**Figure 4 – figure supplement 2 – source data 1. Source data for the quantification of AO-positive SCs.** Source data for the number of AO-positive SCs at 48 hpf depicted in Figure 4 – figure supplement 2B. Data contributed to scatter plot of the number of AO- positive SCs along the pLLN at 48 hpf (mean ± SEM: WT: 0.06 ± 0.04, *cd59^uva48^*: 0.03 ± 0.03; data point = 1 fish). Data was collected from somites 11 to 13 (∼320 µm) and normalized to units per 100 µm.

**Figure 4 – figure supplement 2 – source data 2. Source data for the quantification of AO-positive cells ventral to the pLLN.** Source data for the number of AO-positive cells ventral to the pLLN at 48 hpf depicted in Figure 4 – figure supplement 2C. Data contributed to scatter plot of the number of AO-positive cells ventral to the pLLN at 48 hpf (mean ± SEM: WT: 2.2 ± 0.2, *cd59^uva48^*: 2.8 ± 0.3; dot = 1 fish). Data was collected from somites 11 to 13 (∼320 µm) and normalized to units per 100 µm.

**Figure 4 – figure supplement 2 – source data 3. Source data for the quantification of TUNEL-positive SCs.** Source data for the number of TUNEL-positive SCs at 48 hpf depicted in Figure 4 – figure supplement 2E. Data contributed to scatter plot of the number of TUNEL-positive SCs along the pLLN at 48 hpf (mean ± SEM: WT: 0.4 ± 0.2, *cd59^uva48^*: 0.6 ± 0.3; data point = 1 fish). Data were collected from ten consecutive transverse sections per fish (∼200 µm) and normalized to units per 100 µm.

**Figure 4 – figure supplement 2 – source data 4. Source data for the quantification of TUNEL-positive CNS cells.** Source data for the number of TUNEL-positive CNS cells at 48 hpf depicted in Figure 4 – figure supplement 2F. Data contributed to scatter plot of the number of TUNEL-positive cells in the spinal cord at 48 hpf (mean ± SEM: WT: 0.4 ± 0.2, *cd59^uva48^*: 0.6 ± 0.3; data point = 1 fish). Data were collected from ten consecutive transverse sections per fish (∼200 µm) and normalized to units per 100 µm.

**Figure 5 – source data 1. Source data for quantification of myelin wraps.** Source data for the number of myelin wraps per axon in pLLNs at 7 dpf depicted in Figure 5D. Data contributed to frequency distribution (right) and box and whisker plot (left) of the number of myelin wrappings per pLLN axon at 7 dpf. Data was collected from three sections per fish separated by 100 µm (mean ± SEM: WT: 3.89 ± 0.1, *cd59^uva48^*: 3.08 ± 0.2; p < 0.0001; data point = 1 myelinated axon; n = 3 fish per group).

**Figure 5 – source data 2. Source data for quantification of Na_V_ channel clusters.** Source data for number of Na_V_ channel clusters on the pLLN at 7 dpf depicted in Figure 5E. Data contributed to scatter plot of the number of Na_V_ channel clusters along *mbpa:tagrfp*-positive pLLN nerves at 7 dpf (mean ± SEM: WT: 17.3 ± 0.7, *cd59^uva48^*: 9.9 ± 0.9; p < 0.0001; data point = 1 fish). Data were collected from somites 3 to 13 (∼320 µm) and normalized to units per 100 µm.

**Figure 5 – source data 3. Source data for quantification of NF186 clusters.** Source data for the number of NF186 clusters along the pLLN at 7 dpf depicted in Figure 5F. Data contributed to scatter plot of the number of NF186 clusters along *mbpa:tagrfp*- positive pLLN nerves at 7 dpf (mean ± SEM: WT: 24.0 ± 0.7, *cd59^uva48^*: 18.2 ± 1.3; p = 0.0011; data point = 1 fish). Data were collected from somites 3 to 13 (∼320 µm) and normalized to units per 100 µm.

**Figure 5 – figure supplement 1 – source data 1. Source data for quantification of myelin sheath length.** Source data for myelin sheath measurements depicted in Figure 5 – figure supplement 1D. Data contributed to scatter plot of the length of myelin sheaths (µm) along the pLLN at 7 dpf (mean ± SEM: WT: 66.3 ± 2.4, *cd59^uva48^*: 63.4 ± 4.6; data point = 1 myelin sheath; n = 6 to 7 fish per group).

**Figure 5 – figure supplement 1 – source data 2. Source data for quantification of myelinated nerve volume.** Source data for myelinated nerve volume measurements depicted in Figure 5 – figure supplement 1E. Data contributed to scatter plot of the myelinated nerve volumes (µm^3^) along the pLLN at 7 dpf (mean ± SEM: WT: 5.3 ± 0.3, *cd59^uva48^*: 3.3 ± 0.2; p < 0.0001; data point = 1 fish). Data were collected from somite 12 (∼110 µm). All data were normalized to units per 100 µm.

**Figure 5 – figure supplement 1 – source data 3. Source data for quantification of axon volume.** Source data for axon volume measurements depicted in Figure 5 – figure supplement 1F. Data contributed to scatter plot of axon volumes (µm^3^) along the pLLN at 7 dpf (mean ± SEM: WT: 3.3 ± 0.3, *cd59^uva48^*: 3.3 ± 0.5; data point = 1 fish). Data were collected from somite 12 (∼110 µm). All data were normalized to units per 100 µm.

**Figure 5 – figure supplement 1 – source data 4. Source data for quantification of myelinated axons.** Source data for the number of myelinated axons per pLLN at 7 dpf depicted in Figure 5 – figure supplement 1G. Data contributed to scatter plot (left) of the number of myelinated axons per pLLN at 7 dpf. Data was collected from three sections per fish separated by 100 µm (mean ± SEM: WT: 16.22 ± 0.6, *cd59^uva48^*: 16.67 ± 0.8; data point = 1 section; n = 3 fish per group).

**Figure 6 – source data 1. Source data for quantification of SC-associated MACs.** Source data for MACs-associated with SC membranes at 55 hpf depicted in Figure 6B. Data contributed to scatter plot of the number of MACs in SC membranes at 55 hpf (mean ± SEM: WT: 3.3 ± 0.3, *cd59^uva48^*: 11.6 ± 1.9; p < 0.0001; data point = 1 fish). Data were collected from somites 11 to 13 (∼320 µm) and normalized to units per 100 µm.

**Figure 6 – source data 2. Source data for quantifications of SCs after dexamethasone treatment.** Source data for the number of SCs on the pLLN at 55 hpf after dexamethasone treatment depicted in Figure 6D. Data contributed to scatter plot of the number of pLLN SCs at 55 hpf (mean ± SEM: DMSO: WT: 12.0 ± 0.5, *cd59^uva48^*: 18.1 ± 0.6; Dex: WT: 10.4 ± 0.4, *cd59^uva48^*: 13.4 ± 0.6; p values: WT DMSO vs *cd59^uva48^* DMSO: p <0.0001, WT Dex vs *cd59^uva48^* DMSO: p < 0.0001, WT Dex vs *cd59^uva48^* Dex: p = 0.0014, *cd59^uva48^* DMSO vs *cd59^uva48^* Dex: p < 0.0001; dot = 1 fish). Data were collected from somites 11 to 13 (∼320 µm) and normalized to units per 100 µm.

**Figure 7 – figure supplement 1 – source data 1. Source data for quantification of SCs after extended dexamethasone treatment.** Source data for the number of SCs on the pLLN at 7 dpf after extended dexamethasone treatment depicted in Figure 7 – figure supplement 1B. Data contributed to scatter plot of the number of pLLN SCs at 7 hpf (mean ± SEM: DMSO: 31.9 ± 1.1, Dex: 24.0 ± 0.4; p < 0.0001; data point = 1 fish). Data were collected from somites 3 to 13 (∼320 µm) and normalized to units per 100 µm.

**Figure 7 – source data 1. Source data for quantification of Na_V_ channel clusters after dexamethasone treatment.** Source data for number of Na_V_ channel clusters with dexamethasone treatment depicted in Figure 7C. Data contributed to scatter plot of the number of NaV channel clusters along *mbpa:tagrfp-caax*-positive nerves at 7 dpf (mean ± SEM: DMSO: WT: 26.0 ± 0.8, *cd59^uva48^*: 18.6 ± 0.6; Dex: WT: 23.9 ± 0.8, *cd59^uva48^*: 28.9 ± 0.8; p values: WT DMSO vs *cd59^uva48^* DMSO: p < 0.0001, WT Dex vs *cd59^uva48^* DMSO: p = 0.0001, WT Dex vs *cd59^uva48^* Dex: p = 0.0001, *cd59^uva48^* DMSO vs *cd59^uva48^* Dex: p < 0.0001; data point = 1 fish). Data were collected from somites 3 to 13 (∼320 µm). All data were normalized to units per 100 µm.

**Figure 7 – source data 2. Source data for quantification of myelinated nerve volume after dexamethasone treatment.** Source data for myelinated nerve volume with dexamethasone treatment depicted in Figure 7D. Data contributed to scatter plot of myelinated nerve volumes at 7 dpf (mean ± SEM: DMSO: WT: 5.6 ± 0.4, *cd59^uva48^*: 4.1 ± 0.2; Dex: WT: 4.9 ± 0.1, *cd59^uva48^*: 5.4 ± 0.2; p values: WT DMSO vs *cd59^uva48^* DMSO: p = 0.0009, *cd59^uva48^* DMSO vs *cd59^uva48^* Dex: p = 0.0006; data point = 1 fish). Data were collected from somite 12 (∼110 µm). All data were normalized to units per 100 µm.

---

**Figure 1—Figure Supplement 1.**
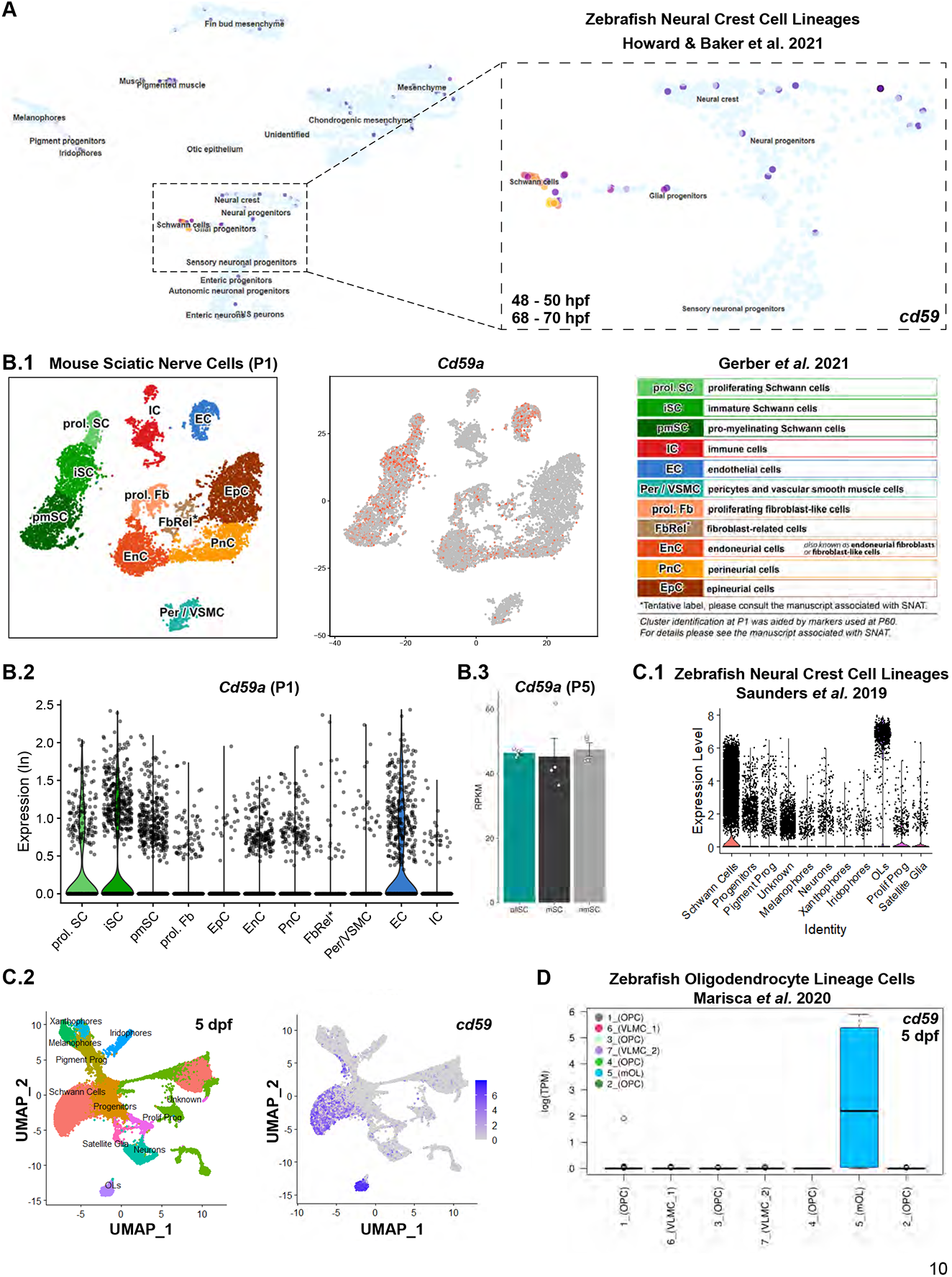
Myelinating glial cell *cd59/Cd59a* expression from bulk and scRNAseq. (**A**) UMAP of *cd59* expression in developing, zebrafish NCC lineages (48 to 50 hpf and 68 to 70 hpf) (Howard et al., 2021). (**B.1**) t-SNE plot and (**B.2**) violin plot of *Cd59a* expression [natural log(ln)] in developing, mouse sciatic nerve cells [postnatal day 1 (P1)] (Gerber et al., 2021). (**B.3**) Bar chart of *Cd59a* expression (RPKM) in developing, mouse sciatic nerve cells (P5) (Gerber et al., 2021). (**C.1**) Violin plot and (**C.2**) UMAP of *cd59* expression in developing, zebrafish NCC lineages (5 dpf) (Saunders et al., 2019). (**D**) Bar chart of *cd59a* expression [log(TPM] in zebrafish, mouse OLCs (5 dpf) (Marisca et al., 2020). All panels were prepared with applications included in the noted publications except for (**C.1 and C.2**), which were assembled by our lab.

**Figure 2—Figure Supplement 1.**
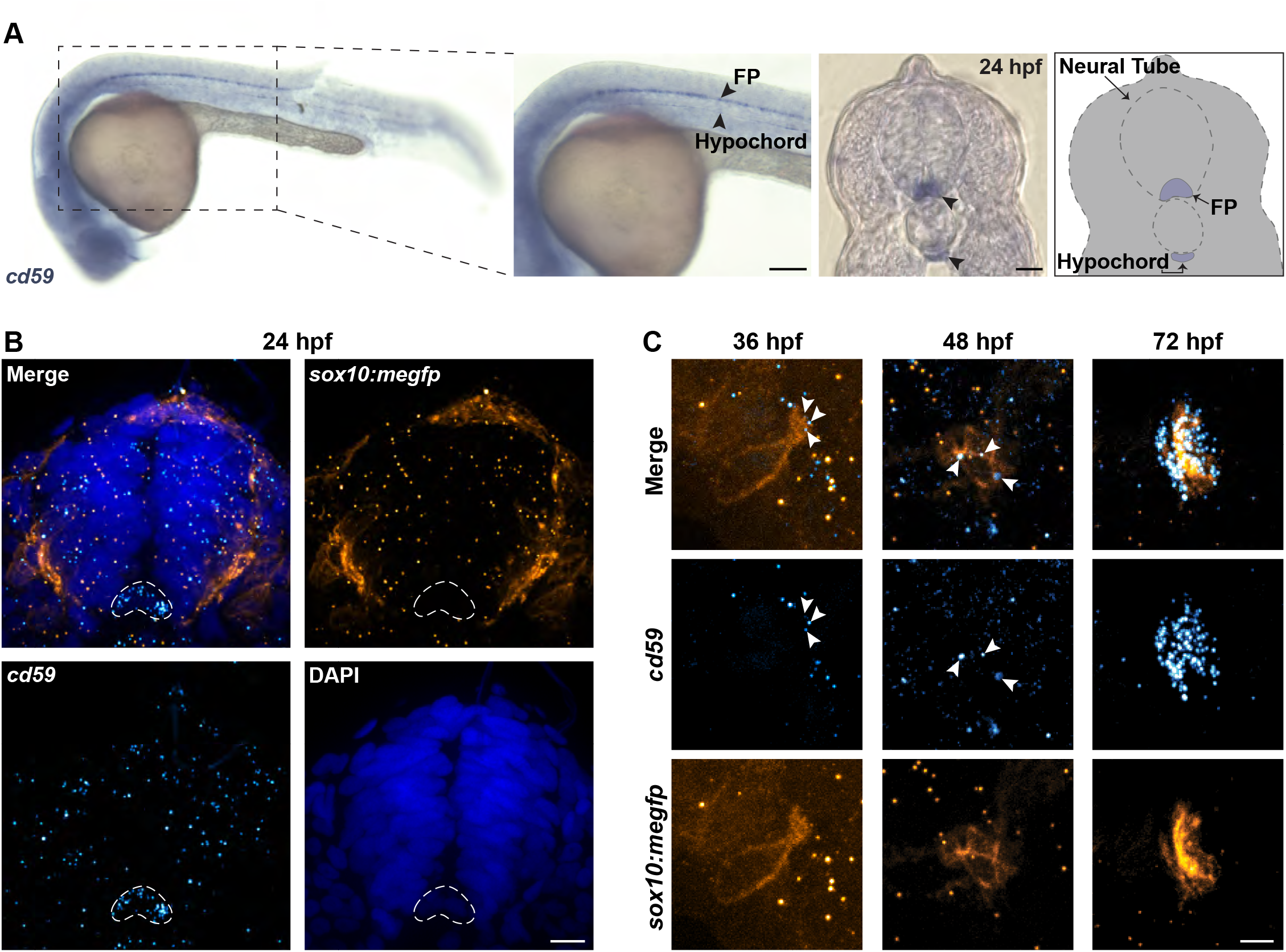
***cd59* is expressed in the floorplate, hypochord, and developing SCs.** (**A**) Whole mount CISH showing *cd59* expression (purple) in the floorplate (FP) and hypochord at 24 hpf. Schematic (right panel) indicates location of *cd59* expression (purple) in FP, hypochord, and neural tube in transverse section. (**B**) FISH showing *cd59* expression (cyan) in the FP (outlined in white dashed lines) and lack of expression in *sox10:megfp*-positive NCCs (orange) in transverse sections. DAPI-positive nuclei indicated in blue. (**C**) FISH showing *cd59* expression (cyan) in *sox10:megfp*- positive (orange) SCPs at 36 hpf, iSCs at 48 hpf, and SCs at 72 hpf on the pLLN. Scale bars: (**A**) lateral view, 100 µm; transverse section, 25 µm; (**B**) 10 µm; (**C**) 5 µm. Artwork created by Ashtyn T. Wiltbank with Illustrator (Adobe).

**Figure 2—Figure Supplement 2.**
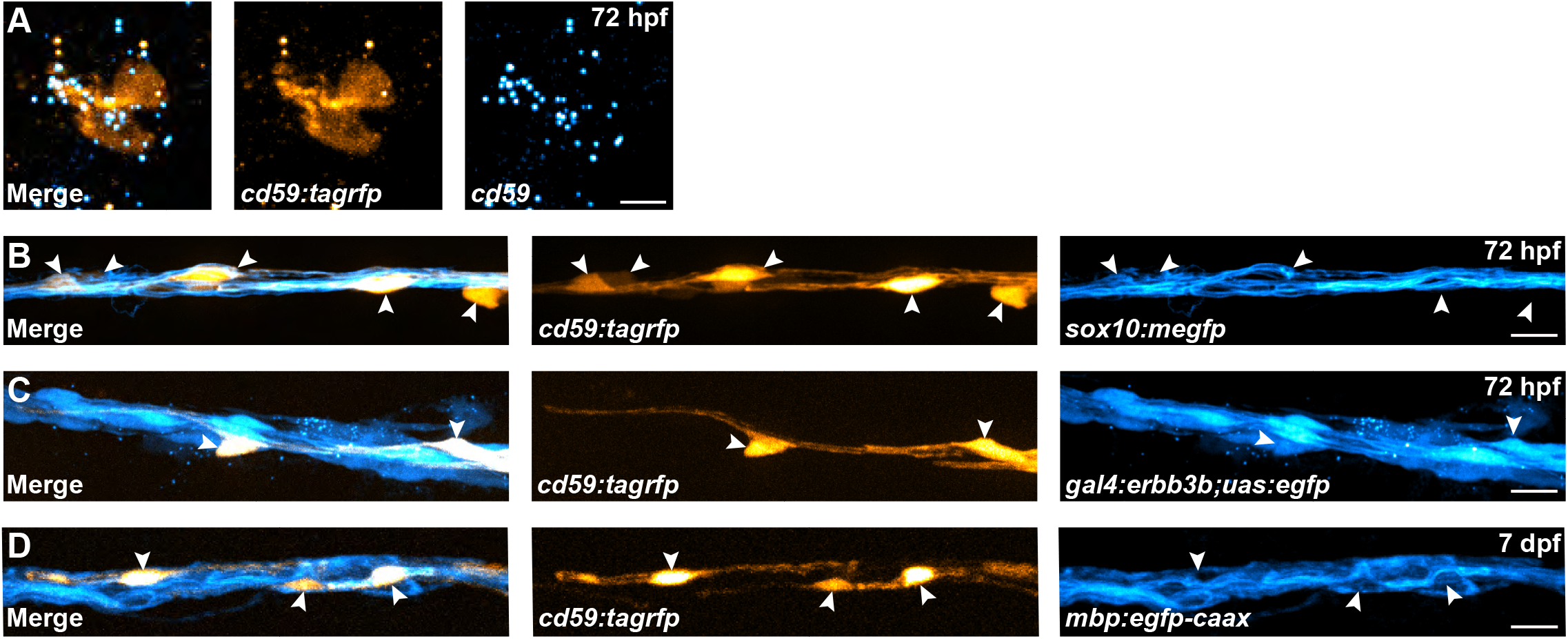
***cd59:tagrfp*-positive SCs express canonical SC markers.** (**A**) FISH showing *cd59* expression (cyan) in *cd59:tagrfp*-positive SC (orange) along the pLLN at 72 hpf. (**B**) *In vivo* imaging showing *cd59:tagrfp*- (orange), *sox10:megfp*- (cyan) positive SCs on the pLLN at 72 hpf. (**C**) *In vivo* imaging showing *cd59:tagrfp*-(orange), *gal4:errb3b;uas:egfp*- (cyan) positive SCs on the pLLN at 72 hpf. (**D**) *In vivo* imaging showing that *cd59:tagrfp*- (orange), *mbp:egfp-caax*- (cyan) positive SCs on the pLLN at 7 dpf. Scale bars: (**A**) 5 µm; (**B - C**) 10 µm.

**Figure 3—Figure Supplement 1.**
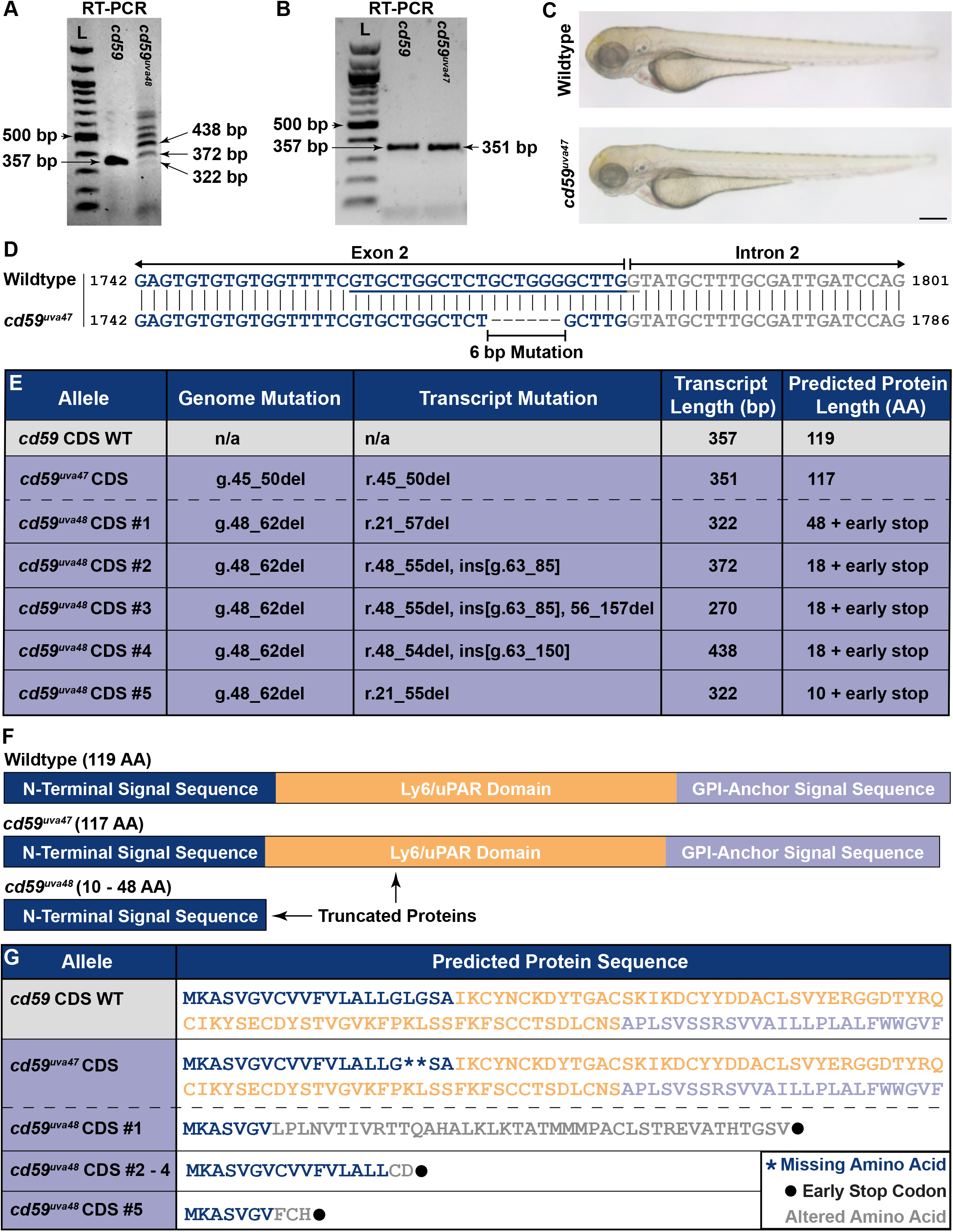
Characterization of *cd59* mutant zebrafish. (**A**) Gel electrophoresis showing wildtype (357 bp) and *cd59^uva48^* (variable transcript size) RT- PCR products at 72 hpf. (**B**) Gel electrophoresis showing wildtype (357 bp) and *cd59^uva47^* (351 bp) RT-PCR products at 72 hpf. RT-PCR products were compared to 100 bp DNA ladder for (**A**) and (**B**). (**C**) Bright-field images of wildtype and *cd59^uva47^* mutant larvae at 72 hpf showing no anatomical defects as a result of *cd59* mutation. (**D**) Genomic sequences for wildtype *cd59* and mutant *cd59^uva47^* showing a 6 bp deletion at the end of exon 2 (blue) in the *cd59* gene. (**E**) Table showing the genome and transcript mutations associated with sequenced RT-PCR products from wildtype (grey) and *cd59* mutant alleles (purple). The resulting transcript and protein length for each allele also included. (**F**) Schematic of Cd59 protein made in wildtypes (119 AA; top panel), *cd59^uva47^* mutants (117 AA; middle panel), and *cd59^uva48^* mutants (10 to 48 AA; middle panel). (**G**) Expasy protein translations (Duvaud et al., 2021) of sequenced RT-PCR products showing altered amino acid sequences (grey text), early stop codons (black dots), and missing amino acids (blue asterisks) detected in *cd59* mutant alleles (purple). Protein domains colored as described previously. Scale bar: (**A**) 0.25 mm. Artwork created by Ashtyn T. Wiltbank with Illustrator (Adobe).

**Figure 4—Figure Supplement 1.**
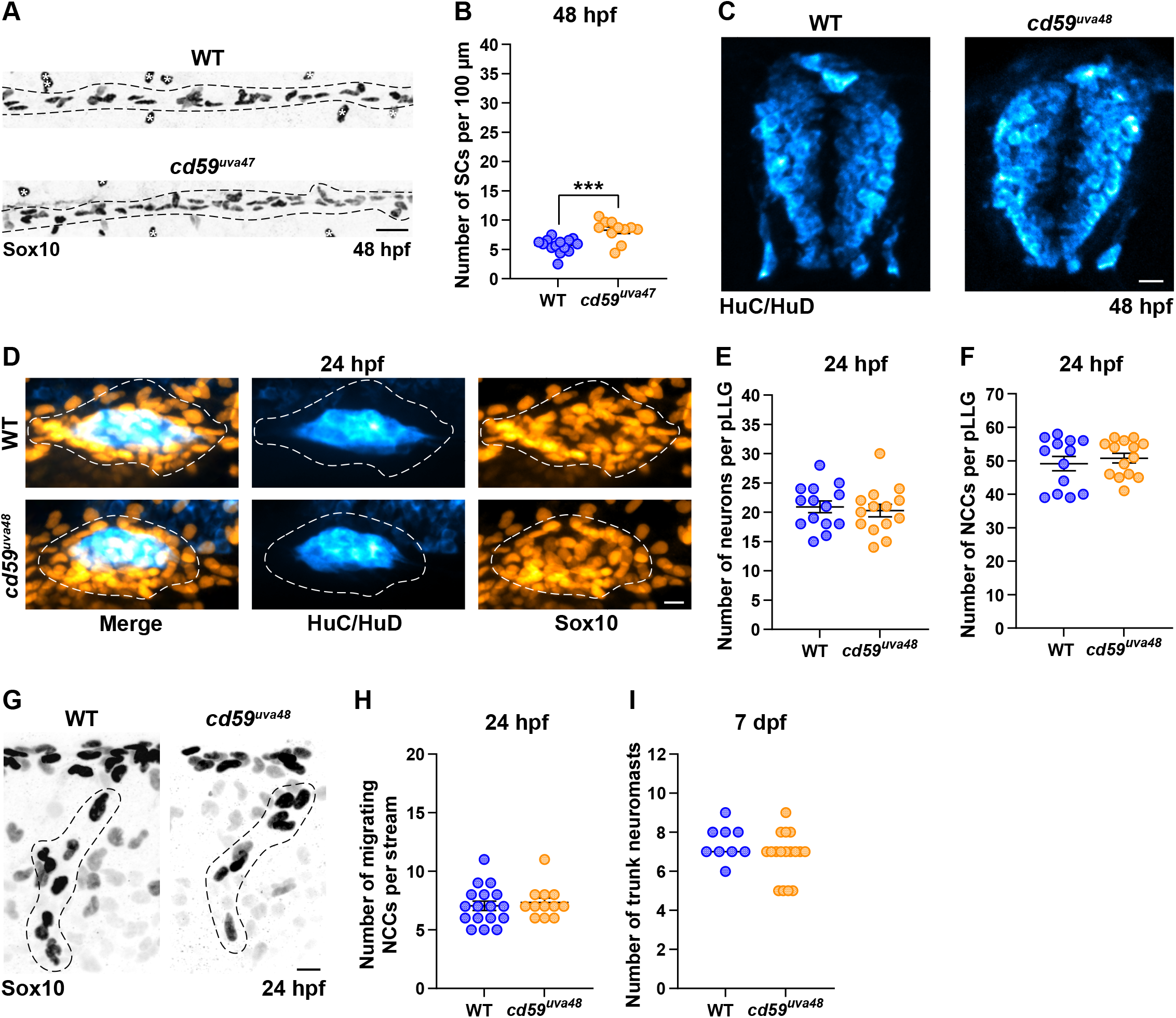
*cd59* does not impact proliferation of neurons or NCCs. (**A**) IF showing Sox10-positive SCs (black/grey) along the pLLN at 48 hpf. Black dashed lines outline the pLLN. Sox10-positive pigment cells outside of the dashed lines (white asterisks) were not included in the analysis. (**B**) Scatter plot of the number of Sox10-positive SCs along the pLLN at 48 hpf (mean ± SEM: WT: 5.7 ± 0.3, *cd59^uva48^*: 8.3 ± 0.5; p = 0.0003; dot = 1 fish). Data was collected from somites 11 to 13 (∼320 µm) and normalized to units per 100 µm. (**C**) IF showing HuC/HuD-positive neurons (cyan) in the spinal cord at 48 hpf (n = 6 fish per group). (**D**) IF showing HuC/HuD-positive neurons (cyan) and Sox10-positive NCCs (orange) in the pLLG at 24 hpf. White dashed lines outline the pLLG. NCCs outside of the dashed lines were not included in the analysis. (**E**) Scatter plot of the number of HuC/HuD-positive neurons in the pLLG at 24 hpf (mean ± SEM: WT: 20.9 ± 1.0, *cd59^uva48^*: 20.3 ± 1.1; dot = 1 fish). (**F**) Scatter plot of the number of Sox10-positive NCCs in the pLLG at 24 hpf (mean ± SEM: WT: 49.2 ± 2.1, *cd59^uva48^*: 50.8 ± 1.4; dot = 1 fish). (**G**) IF showing Sox10-positivie NCCs (black/grey) migrating in the trunk at 24 hpf. Black dashed lines outline the migrating NCCs. NCCs outside of the dashed lines were not included in the analysis. Data collected from the 12^th^ somite. (**H**) Scatter plot of the number of Sox10-positive, migrating NCCs in the trunk at 24 hpf (mean ± SEM: WT: 7.1 ± 0.4, *cd59^uva48^*: 7.3 ± 0.4; dot = 1 fish). (**I**) Scatter plot of the number of Tubulin-positive neuromasts in the trunk at 7 dpf (mean ± SEM: WT: 7.4 ± 0.3, *cd59^uva48^*: 6.8 ± 0.3; dot = 1 fish). Scale bars: (**A**) 25 µm; (**C, D, G**) 10 µm. **Video 1.** Wildtype *sox10:tagrfp*-positive SCs along the pLLN migrate and undergo cell division from 48 to 55 hpf. Images were taken every 10 minutes, and the movie runs at 3 frames per second (fps). Data were collected from somites 11 to 13 (∼320 µm). Mitotic events indicated with magenta arrows. Quantification of the number of mitotic events is indicated in Figure 4H. Scale bar: 25 µm. **Video 2.** *cd59^uva48^* mutant *sox10:tagrfp*-positive SCs along the pLLN migrate and undergo cell division from 48 to 55 hpf. Images were taken every 10 minutes, and the movie runs at 3 frames per second (fps). Data were collected from somites 11 to 13 (∼320 µm). Mitotic events indicated with magenta arrows. Quantification of the number of mitotic events is indicated in Figure 4H. Scale bar: 25 µm. **Video 3.** Wildtype *sox10:tagrfp*-positive SCs along the pLLN migrate, undergo cell division, and begin to form myelin from 54 to 75 hpf. Images were taken every 10 minutes, and the movie runs at 3 fps. Data were collected from somites 11 to 13 (∼320 µm). Mitotic events indicated with magenta arrows. Quantification of the number of mitotic events is indicated in Figure 4I. Time of last final cell division is indicated in Figure 4J. Scale bar: 25 µm. **Video 4.** *cd59^uva48^* mutant *sox10:tagrfp*-positive SCs along the pLLN migrate, undergo cell division, and begin to form myelin from 54 to 75 hpf. Images were taken every 10 minutes, and the movie runs at 3 fps. Data were collected from somites 11 to 13 (∼320 µm). Mitotic events indicated with magenta arrows. Quantification of the number of mitotic events is indicated in Figure 4I. Time of last final cell division is indicated in Figure 4J. Scale bar: 25 µm.

**Figure 4—Figure Supplement 2.**
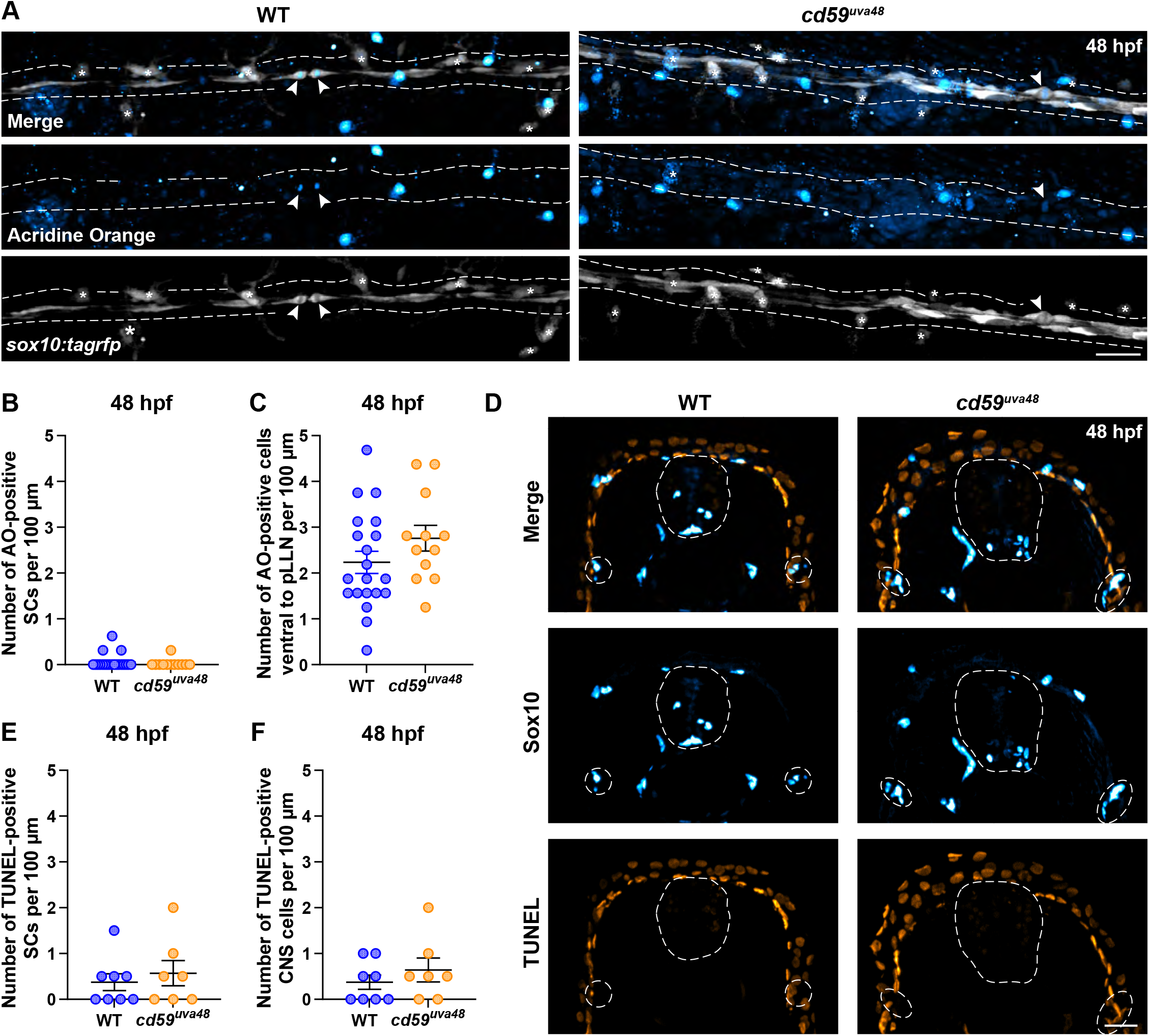
Loss of *cd59* does not impact cell death within the PNS and CNS. (**A**) AO incorporation assay showing AO-positive (cyan), *sox10:tagrfp*- positive SCs (grey) along the pLLN at 48 hpf. White dashed lines outline the pLLN. Arrowheads denote proliferative events, which can also be detected by AO. (**B**) Scatter plot of the number of AO-positive SCs along the pLLN at 48 hpf (mean ± SEM: WT: 0.06 ± 0.04, *cd59^uva48^*: 0.03 ± 0.03; dot = 1 fish). (**C**) Scatter plot of the number of AO-positive cells ventral to the pLLN at 48 hpf (mean ± SEM: WT: 2.2 ± 0.2, *cd59^uva48^*: 2.8 ± 0.3; dot = 1 fish). Data was collected from somites 11 to 13 (∼320 µm) and normalized to units per 100 µm for (**B**) and (**C**). (**D**) TUNEL assay showing TUNEL-positive (orange) cells in the spinal cord and Sox10-positive SCs along the pLLN at 48 hpf. White dashed lines outline the spinal cord (medial) and the pLLNs (lateral). (**E**) Scatter plot of the number of TUNEL-positive SCs along the pLLN at 48 hpf (mean ± SEM: WT: 0.4 ± 0.2, *cd59^uva48^*: 0.6 ± 0.3; dot = 1 fish). (**F**) Scatter plot of the number of TUNEL-positive cells in the spinal cord at 48 hpf (mean ± SEM: WT: 0.4 ± 0.2, *cd59^uva48^*: 0.6 ± 0.3; dot = 1 fish). Data were collected from ten consecutive transverse sections per fish (∼200 µm) and normalized to units per 100 µm for (**E**) and (**F**). Scale bars: (**A, D**) 25 µm.

**Figure 5—Figure Supplement 1.**
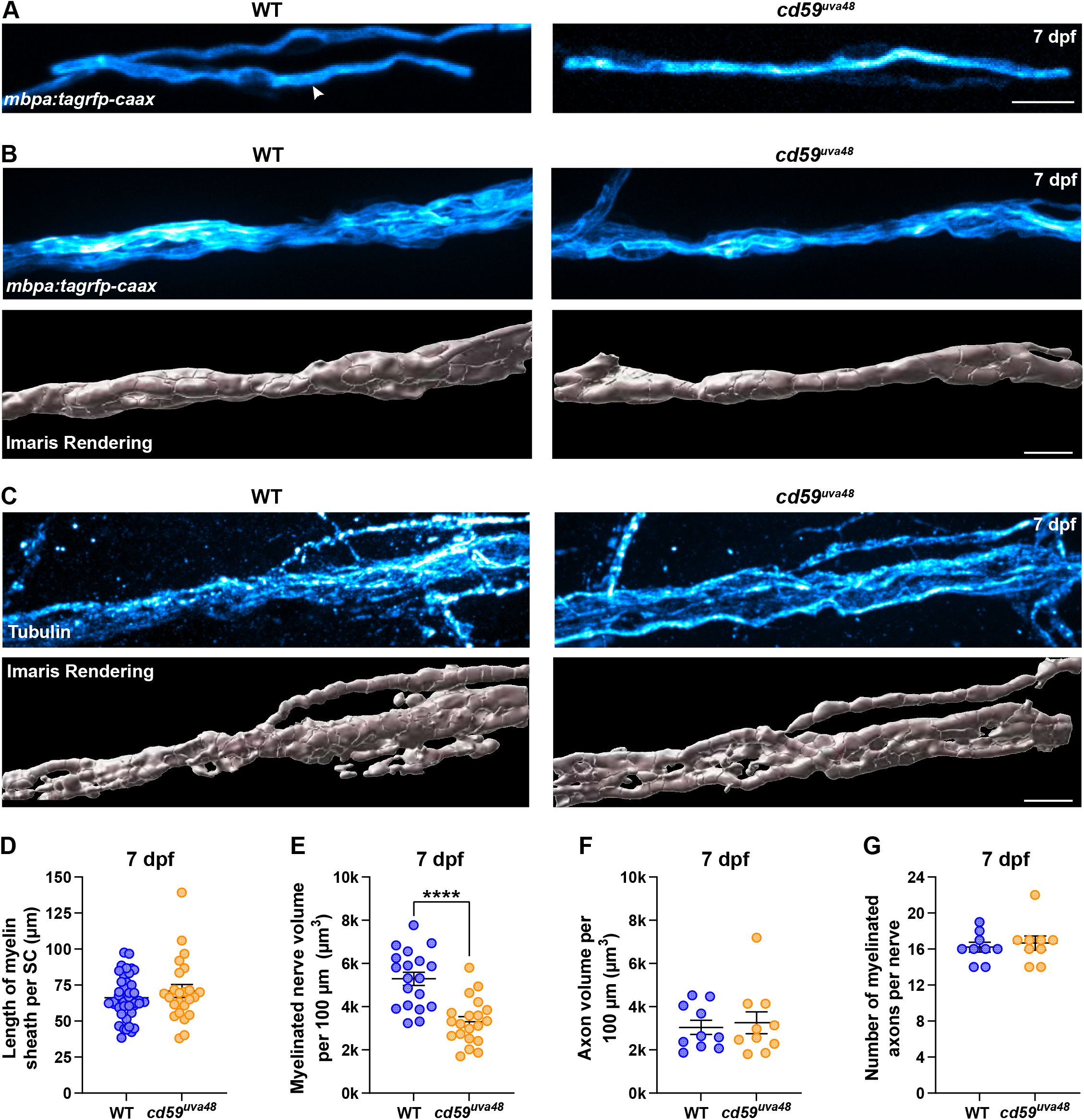
Myelin volume is reduced in *cd59^uva48^* mutants. (**A**) Mosaic labeling of *mbpa:tagrfp-caax*-positive SC myelin sheaths (cyan) along the pLLN at 7 dpf. (**B**) *In vivo* imaging of *mbpa:tagrfp-caax*-positive pLLN nerves (cyan; top panel) at 7 dpf. Imaris renderings (white) of nerve volume in bottom panel. (**C**) IF showing Tubulin-positive pLLN axons (cyan; top panel) at 7 dpf. Imaris renderings (white) of axon volume in bottom panel. Representative images for (**B**) and (**C**) are from somite 12 (∼110 µm). (**D**) Scatter plot of the length of myelin sheaths (µm) along the pLLN at 7 dpf (mean ± SEM: WT: 66.3 ± 2.4, *cd59^uva48^*: 63.4 ± 4.6; dot = 1 myelin sheath; n = 6 to 7 fish per group). (**E**) Scatter plot of the myelinated nerve volumes (µm^3^) along the pLLN at 7 dpf (mean ± SEM: WT: 5.3 ± 0.3, *cd59^uva48^*: 3.3 ± 0.2; p < 0.0001; dot = 1 fish). (**F**) Scatter plot of axon volumes (µm^3^) along the pLLN at 7 dpf (mean ± SEM: WT: 3.3 ± 0.3, *cd59^uva48^*: 3.3 ± 0.5; dot = 1 fish). (**G**) Scatter plot (left) of the number of myelinated axons per pLLN at 7 dpf. Data was collected from three sections per fish separated by 100 µm (mean ± SEM: WT: 16.22 ± 0.6, *cd59^uva48^*: 16.67 ± 0.8; dot = 1 section; n = 3 fish per group). Data quantified in (**G**) were determined from electron micrographs in (Figure 5A). Scale bars: (**A** - **C**) 10 µm.

**Figure 6—Figure Supplement 1.**
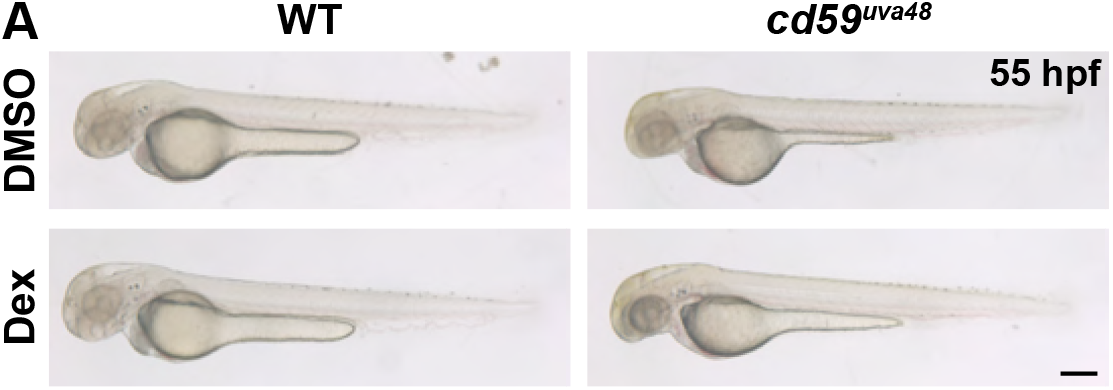
Dexamethasone does not interfere with development. (**A**) Bright-field images of wildtype and mutant embryos (55 hpf) treated with DMSO or 100 µM Dex from 24 to 55 hpf. Scale bar: 0.25 mm.

**Figure 7—Figure Supplement 1.**
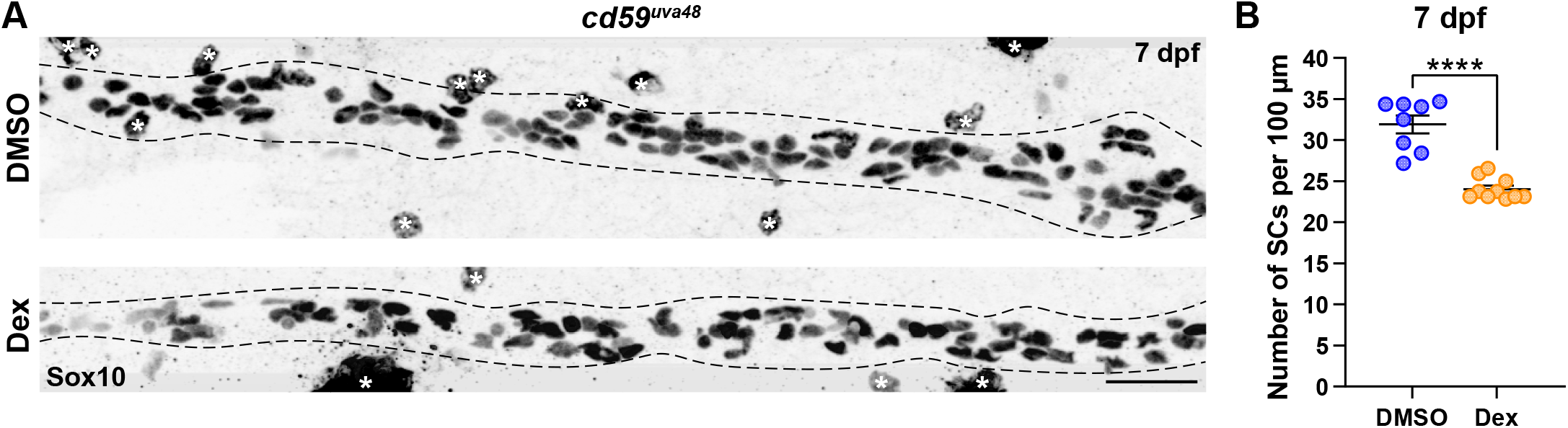
Extended dexamethasone treatment had same impact on SC proliferation in *cd59^uva48^* mutant larvae. (**A**) IF showing Sox10-positive SCs (black/grey) at 7 dpf in *cd59^uva48^* mutant larvae treated with DMSO or 100 µM Dex from 24 to 75 hpf. Black dashed lines outline the pLLN. Sox10-positive pigment cells outside of the dashed lines (white asterisks) were not included in the analysis. (**B**) Scatter plot of the number of pLLN SCs at 7 hpf (mean ± SEM: DMSO: 31.9 ± 1.1, Dex: 24.0 ± 0.4; p < 0.0001; dot = 1 fish). Data were collected from somites 3 to 13 (∼320 µm) and normalized to units per 100 µm. Scale bar: 25 µm.

## Footnotes

* According to the myelin length data noted in Figure 5 – figure supplement 1D, most myelin sheaths are less than 100 µm. Separating each section by 100 µm would therefore enable quantification of three separate groups of SCs per pLLN.

